# A distinct subset of stem-cell memory is poised for the cytotoxicity program in CD4^+^ T cells in humans

**DOI:** 10.1101/2025.04.29.651205

**Authors:** Raunak Kar, Shreya Sinha, Zainab Khatun, Anjali Sharma, Veena S. Patil

## Abstract

The peripheral CD4^+^ T-cells with cytolytic potential (CD4-CTLs) have been detected in several diseases, including infectious diseases and various cancers. They form the component of protective immune response and are shown to be enriched in the effector memory expressing CD45RA (T_EMRA_) in humans. However, the lack of understanding about their lineage, molecular characters, and cytolytic potential in comparison to CD8-CTLs has restricted their utility. Thus, in this study, by parallelly analysing the CD4-CTLs and CD8-CTLs, we demonstrate that they are indistinguishable for the cytolytic program, with both showing similar gene expression profile and T cell antigen-receptor (TCR) clonal expansion in humans. Further, using an integrative multi-omics analysis combining the transcriptome, TCR repertoire, and open chromatin profile of CD4^+^ naïve (CD4-T_N_) and memory T-cell subsets, we discovered a distinct stem-cell memory subset (T_SCM-CTL_) that is pre-committed to cytolytic program that shared significant TCR clonotypes with expanded CD4-CTL-effectors. Further, through an *in vitro* differentiation model, we developed CD4^+^ T-cells with cytolytic potential (iCD4-CTLs) from CD4-T_N_ cells that showed gradual and progressive acquisition of cytolytic program, exhibiting progressive chromatin accessibility at cytotoxicity-associated genes. Of particular interest was the property of iCD4-CTLs that co-expressed longevity-associated genes along with cytotoxicity-associated genes, hence generating long-lived CD4-CTL-effectors. Overall, the process of generation of iCD4-CTLs from CD4-T_N_, deciphered the molecular signatures of early commitment to the cytotoxicity program. Together, our study advocates for exploring both CD4-CTLs and CD8-CTLs for vaccine development, vaccine efficacy testing, as well as immunotherapies and cell-based therapies for precision medicine.

**Teaser:** Integrative multi-omics analysis of CD4^+^ T cell subsets revealed the cytolytic potential of CD4-T_EMRA_ cells in comparison to CD8-T_EMRA_ and their origin from stem-cell memory.

## INTRODUCTION

The naïve T cells activated during the primary infection, in addition to differentiating to effector cells, also forms a pool of memory T cells that have the potential to elicit a quicker and a stronger immune response against the same pathogen during a secondary infection. The T cell memory pool is heterogeneous and can be further classified based on longevity, proliferative capacity and effector-ness. Early studies described memory as either long-lived T central memory (T_CM_) or short-lived effector memory (T_EM_) and T_EM_ expressing CD45RA (T_EMRA_) based on the surface expression of cell adhesion molecule, L-selectin (CD62L); isoform of tyrosin phosphatase CD45, CD45RA; co-stimulatory molecule, CD27; and chemokine receptor type 7, CCR7; within the CD4^+^ and CD8^+^ T cell subsets in humans (*1–3*). Many years later the antigen-experienced stem like memory (T_SCM_) was discovered within the naïve compartment based on the surface expression of CD95 and IL2RA (*4, 5*). These memory subsets define developmental stages in the memory formation and are better described for CD8^+^ T cell subset where a more linear development from naïve -> T_SCM_ -> T_CM_ -> T_EM_ -> T_EMRA_ has been suggested (*5*). Further the CD4^+^ T helper (T_H_) memory cells can be classified based on their functional properties into T_H_1, T_H_2, T_H_17, T_H_1/17, T follicular helper (T_FH_) and T regulatory cells (T_REGs_), where each of them have a specialized function (*6–9*). T-bet expressing T_H_1 cells that secrete interferon gamma (IFNγ) are known for their role in viral infections, BCL-6 expressing CD4^+^ T follicular helper (T_FH_) cells can secrete IL21 and assist B cells (*6–8, 10*). The CD4^+^ helper T cells can also be regulators of immune response via the T regulatory cells (T_REGs_) that express FOXP3 and secrete TGFβ and IL10 cytokines (*11*). Though the functional CD4^+^ T_H_ memory subsets are relatively well defined, comprehensive studies describing the developmental memory subsets for their lineage, T cell antigen-receptor (TCR) clonal expansion and epigenetic program are far fewer (*12–14*). Each T_H_ memory subset can exist in both long-lived and effector memory subsets suggesting their different developmental stages (*7*).

Though classically the CD4^+^ T cells are known for their helper roles and the CD8^+^ T cells for their cytotoxic roles in immune responses, the Major Histocompatibility Complex (MHC)II restricted CD4^+^ T cells have also been shown to be killers in a wide array of infectious diseases, cancers and autoimmune disorders for many decades now (*15–21*). The circulating CD4^+^ cytotoxic T lymphocytes (CD4-CTLs) have been better studied in viral infections in both humans and animal models, which include human cytomegalovirus (hCMV), Epstein-Barr virus (EBV), human immunodeficiency virus (HIV), dengue virus (DENV), Influenza virus (IAV), severe acute respiratory syndrome coronavirus 2 (SARS-CoV2) and many more (*12, 17, 18, 22–30*). Most importantly, the magnitude of the CD4-CTL response has been associated with better clinical outcomes in both acute and chronic viral infections and anti-tumor immune responses (*31*). Successful vaccinations against yellow fever virus (YFV), IAV, smallpox, vaccinia virus, poliovirus and HIV have also been shown to elicit CD4-CTL responses (*31–33*). Thus, eliciting a strong CD4-CTL response has been considered an important goal of vaccination against several viral infections.

The CD4-T_EMRA_ memory compartment has been shown to be enriched for CD4-CTLs (*12, 13, 30*). However, the molecular and epigenetic landscapes that drive the differentiation, maintenance, and function of human CD4-CTLs from naïve T cells are still elusive. Further, though it has been shown that CD4-T_EMRA_ expresses cytotoxicity-associated genes and show enrichment for the overall cytotoxic program, they have not been well studied in conjunction with the bona fide cytotoxic T cells, the CD8-CTLs (CD8-T_EMRA_) to assess their cytolytic potential (*12*). Hence, here, through global transcriptomic- and cytotoxicity-related protein expression analysis from the same donors, we show that CD4-CTLs and CD8-CTLs are indistinguishable for cytotoxicity program and both show increased TCR clonal expansion, the mark of the effector cells. Next, using both bulk and single-cell transcriptomic analysis paired with TCR analysis, we identified T_SCM-CTLs_, stem-like memory cells poised for CD4-CTL lineage within the CD4-T_SCM_ subset. Further, through an *in vitro* differentiation model, we systematically developed CD4 T cells with cytolytic ability from CD4 naïve T-cells, established a differentiation pathway and identified transcriptomic as well epigenomic regulators of the CTL program development that overlaps with *ex vivo* CD4-T_EMRA_ effector cells, thus establishing the CD4-CTL lineage from naïve cells via the T_SCM-CTLs_.

## RESULTS

### The cytolytic program of CD4-CTLs and CD8-CTLs is indistinguishable

The CD4^+^ cytotoxic T lymphocytes (CD4-CTLs) have been shown to be enriched in CD4-T_EMRA_ (effector memory expressing CD45RA; CD4^+^CD45RA^+^CCR7^-^) memory compartment which was further found to be heterogenous with mixtures of effectors and precursors of CD4-CTLs, that could be distinguished based on IL7R expression (*12, 13*). The proportion of CD4-T_EMRA_ varies significantly across individuals and is positively correlated with better clinical outcome in viral infections and vaccination (*12, 13, 28, 30, 33*). Consistent with previous report, we observed a huge variability in the CD4-T_EMRA_ proportion, ranging from 0.11% to over 25% of the total CD4^+^ T cells in the peripheral blood across 110 subjects (Fig. 1, A and B) (*12*). Further, we noted a significant positive correlation between frequency of CD4-T_EMRA_ and the proportion CD4-T_EMRA_ expressing cytolytic molecules such as granzyme B (GZMB), CD244 (2B4), CX3CR1, KLRG1 and GPR56 (ADGRG1) (Fig. 1C and table S1) (*12, 13, 30, 31*). Similarly, upon stimulation, the CD4-T_EMRA_ cells produced interferon gamma (IFNγ), a cytokine made by cytotoxic cells, and expressed the degranulation marker, LAMP1, on the cell surface, further highlighting the cytolytic ability of these cells (Fig. 1D, and fig. S1A) (*12, 13, 30, 34*). Hence, to identify and understand the overall cytotoxic program of CD4-CTLs in relation to the other CD4^+^ memory and naïve T cell subsets we performed RNA-Sequencing (RNA-Seq) of *ex vivo* isolated CD4^+^ T cell memory compartments; CD4-T_N_ (Naïve T cells; CD4^+^CD45RA^+^CCR7^+^CD95^-^), CD4-T_SCM_ (stem cell memory; CD4^+^CD45RA^+^CCR7^+^CD95^+^), CD4-T_CM_ (central memory; CD4^+^CD45RA^-^CCR7^+^), CD4-T_EM_ (effector memory; CD4^+^CD45RA^-^CCR7^-^), CD4-CD127^hi^ T_EMRA_ (precursor CD4-CTL or CD4-T_EMRA-P_), and CD127^lo^ CD4-T_EMRA_ (effector CD4-CTL or CD4-T_EMRA-E_) from 10 healthy human subjects with higher CD4-T_EMRA_ proportion (Fig. 1, A, E and F, and fig. S1, B to D) (*1, 4, 5, 12*). The transcriptomic data analysis using the 2000 most variable transcripts, showed that the T_SCM_ and T_CM_ subsets resembled that of naïve T cells and showed enrichment for signatures of longevity, while the T_EM_ and the precursor- and effector-T_EMRA_ subsets clustered together, and showed enrichment for effector molecules (*GZMB*, *PRF1*, *CD244*, *CX3CR1*, *FCGR3A*, *IFNG* etc) (Fig. 1, E and F, and fig. S1, C and D). We then compared the transcriptomes of the CD4-CTLs (CD4-T_EMRA-P_ and CD4-T_EMRA-E_) with CD8-CTLs (CD8^+^-T_EMRA_) from the same donors. We observed that they clustered together in a principal component analysis (PCA) and expressed cytotoxicity-associated transcripts at comparable levels, indicating shared transcriptomic signatures between CD4-CTLs and CD8-CTLs (Fig. 1, A, E and F). Importantly, even the pairwise comparison of CD4-T_EMRA_ and the CD8-T_EMRA_ subsets did not identify any notable differences in the expression of transcripts associated with the cytotoxicity program (Fig. 1, G and H, and fig. S1E, and data file S1). The expression of cytolytic molecules such as *GZMB*, *GNLY*, *PRF1*, *ADGRG1 (GPR56)*, *KLRG1*, *CX3CR1*, and transcription factors (TFs) and the long non-coding RNA (lncRNA) associated with CTLs, *TBX21* (encoding T-bet) and *linc02384* respectively, was comparable between CD4-CTLs and CD8-CTLs (Fig. 1H, and data file S1) (*12*). Further, the gene set enrichment analysis (GSEA) using the known gene sets enriched in CD8-effectors, CD4-CTL-effectors and CD4-T_EMRA_ showed no significant differences between CD4-CTLs and CD8-CTLs (Fig. 1I, and data file S2) (*12, 35, 36*). Next, to assess if the differences could be captured at the protein level, we performed flow cytometry analysis to examine the expression of various proteins and TF associated with cytotoxicity (GNLY, GZMB, PRF1, GPR56 (ADGRG1), CX3CR1, KLRG1, and T-bet) between CD4-CTLs and CD8-CTLs (Fig. 1J). Even at protein level, most of these CTL associated molecules showed no significant differences between CD4-T_EMRA_ and CD8-T_EMRA_ (Fig. 1, J and K). Overall, these results show that there are no notable differences between CD4-CTL and CD8-CTL memory subsets, and both share similar gene expression profiles.

**Fig. 1.**
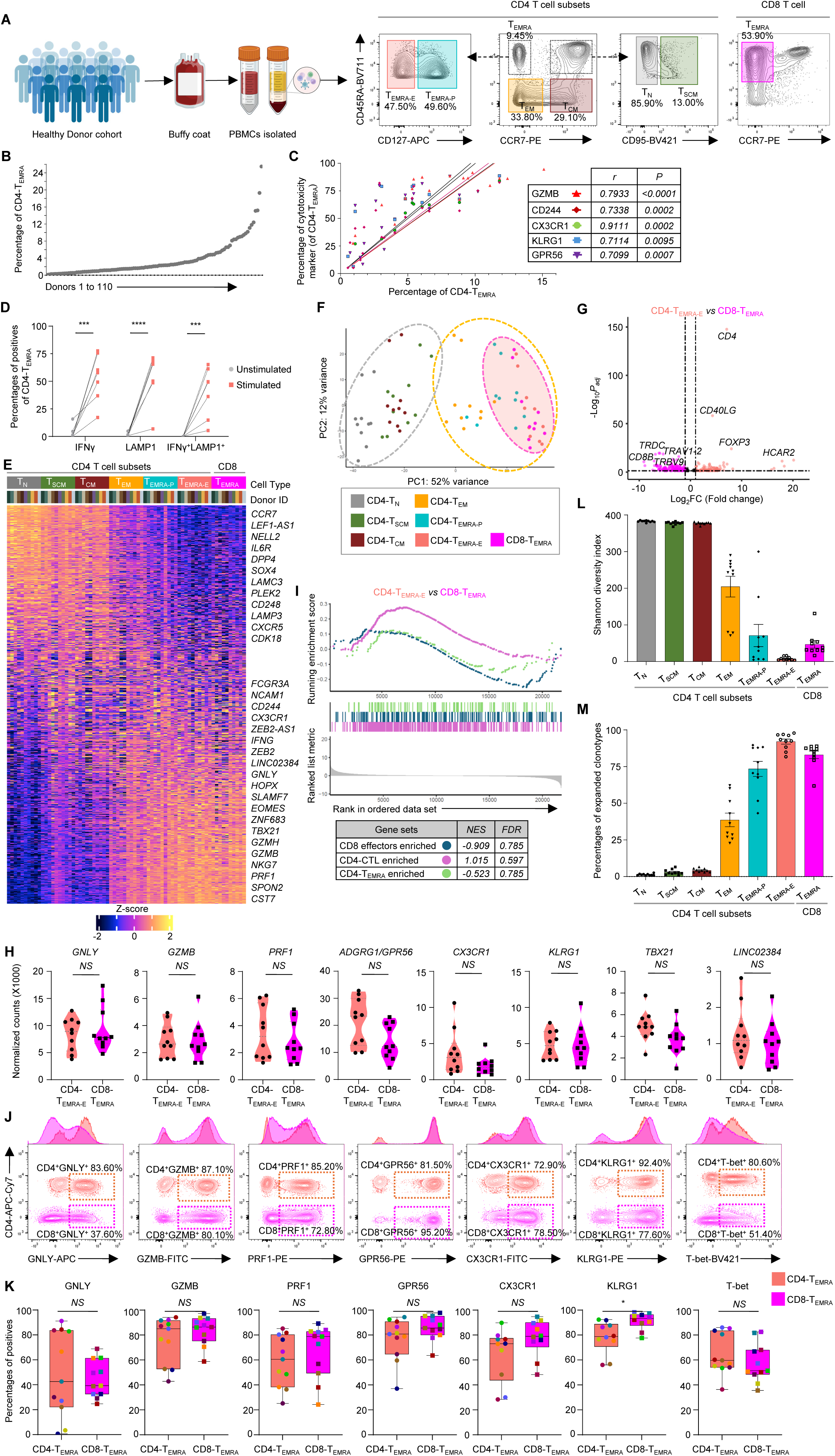
The CD4-T_EMRA_ subset resembles the CD8-T_EMRA_ subset. (A) Schematic representation of study design employed for isolation of memory T cell subsets in humans using flow cytometry. The percentages in the contour plots are of parent population. (B) Scatter plot shows the proportions of CD4-T_EMRA_ of CD4^+^ T cells in PBMCs of 110 healthy human donors. (C) Scatter plot of Pearson correlation between the proportion of CD4-T_EMRA_ and the proportion of CD4-T_EMRA_ positive for the indicated cytotoxicity-related molecules. Each colored line represents the linear regression of correlation with 95% confidence interval of indicated marker, where each colored point represents a donor (n=16-20). The correlation *r* value and *P* value from Student’s paired two-tailed T test for each molecule is tabulated. (D) Dot plot shows the percentage of IFNγ^+^, LAMP1^+^ and IFNγ^+^LAMP1^+^ cells in CD4-T_EMRA_ compartment with and without stimulation. Each dotted line connects the same donor (n=7). (E) Heatmap of bulk transcriptomic analysis shows the row-wise z-score of normalized counts of 2000 most variable genes across the indicated cell type from 10 donors. (F) Principal component analysis (PCA) plot for 2000 most variable genes across the T cell compartments from 10 donors. (G) Volcano plot for differentially expressed transcripts (based on DESeq2) between CD4-T_EMRA-E_ (peach) and CD8-T_EMRA_ (pink), where the genes are colored based on cell type where they are upregulated. (H) Violin plots show the normalized counts for the indicated cytotoxicity-related transcripts in CD4-T_EMRA-E_ and CD8-T_EMRA_. *NS*=non-significant based on DESeq2. (I) Combined GSEA plot for indicated gene sets comparing CD4-T_EMRA-E_ and CD8-T_EMRA_. Individual normalized enrichment scores (NES) and false discovery rates (FDR) are tabulated. (J) Representative contour plots show the expression of the indicated proteins within the CD4-T_EMRA_ (peach) and CD8-T_EMRA_ (pink) compartment in a flow-cytometry analysis. Adjacent histograms show geometric mean fluorescence intensity (MFI). Percentages are of parent population. (K) Box plots with median and interquartile range show the percentage of positives for the indicated proteins within the CD4-T_EMRA_ and CD8-T_EMRA_ compartment in a flow-cytometry analysis from 9-11 donors. Each dot represents a donor. \**P* <0.05, *NS P*>0.05 from Student’s paired two-tailed T test. (L-M) Bar graphs show the mean Shannon-Weiner diversity index (L) and proportions of expanded clonotypes (≥3) (M) for TCRβ clonotypes n=10 donors.

The T cells are known for the diversity in the T cell antigen receptor (TCR) clonotype with every cell having a unique clonotype unless they have resulted from clonal expansion in response to an infection or an immunological event, hence effector cells are known to show restricted TCR repertoire (*37*). Thus, to understand the TCR clonal diversity and clonal expansion across the memory subsets, we performed TCR-Seq analysis of both TCRα and TCRβ chains across CD4^+^ naïve and memory subsets and CD8-T_EMRA_ (*38, 39*). TCR repertoire analysis showed that similar to naïve T cells, the long-term memory subsets T_SCM_ and T_CM_ have highly diverse TCRα and TCRβ repertoire, while the effector memory subsets have limited diversity (Fig. 1L, fig. S1F, and data file S3). On the other hand, similar to the CD8-T_EMRA_ subset, the CD4-T_EMRA_ subsets are hugely clonally expanded (Fig. 1M, fig. S1G, and data file S3) (*12*). We further noted preferential usage of a few V and J genes over others that could not be correlated with their genomic locus, in both TCRα and TCRβ clonotypes across the memory subsets highlighting the random selection of V and J genes during recombination at the TCR loci (fig. S1H, and data file S3). The single-cell transcriptomic and TCR repertoire analysis of rare hCMV-specific memory T cells has also revealed similar observations (*29*). Together, these data show that the CD4-CTL and CD8-CTL memory compartments are very similar for their cytotoxicity program and are the result of clonal expansion, indicating that the program employed for cytolysis by these cells is same and the difference potentially comes from how they recognise the pathogen.

### CD4^+^ T stem-cell memory (T_SCM_) shows dual signatures

Considering, the non-classical cytotoxic nature of CD4^+^ T cells in the CD4-T_EMRA_ compartment, we next wanted to delineate the developmental trajectory of CD4-CTLs by identifying a potential long-lived memory subset that is pre-committed to the CTL lineage in the CD4^+^ memory T-cell subsets. To this end, we first compared the expression of transcripts that distinguish CD4-T_N_ and CD4-T_EMRA-E_ across all the developmental CD4^+^ T cell memory subsets (Fig. 2A, and data file S1). Consistent with our previous observation, largely the long-term memory subsets (T_SCM_ and T_CM_) resembled CD4-T_N_ while the effector memory subsets (T_EM_ and T_EMRA-P_) resembled CD4-T_EMRA-E_, and clustered as two groups in PCA analysis (Fig. 1, E and F, 2A, and fig. S2A). The transcripts such as *TCF7*, *SELL*, *FOXP1*, *CCR7*, *NELL2*, *LTB*, *CD27* that were upregulated in CD4-T_N_ compared to T_EMRA-E_, were also expressed at an elevated level by T_SCM_ and T_CM_ when compared to effector memory, although the expressions were further reduced in them compared to CD4-T_N_ (Fig. 2, A and B, and data file S1). The T_SCM_ subset could be further distinguished from CD4-T_N_ based on the expression of *FAS (CD95)*, *CXCR3*, *IL2RB* and *CD58*, which represent the status of antigenic-experience (fig. S2B) (*4, 5*). The T_SCM_ subset was also marked by the highest expression of the stem-cell marker, *CD38* (Fig. 2B) (*40*). Similarly, the transcripts upregulated in effectors (*GZMB*, *PRF1*, *NKG7*) were also expressed by T_EM_ and T_EMRA-P_ subsets albeit at a reduced level compared to T_EMRA-E_ (Fig. 2, A and B) (*12*). However, interestingly we noted a significantly higher expression of the cytotoxicity-associated transcripts (*GZMB*, *NKG7*, *PRF1*, *ADGRG1*, *CD244*, *ZEB2*, *ZNF683*) in T_SCM_ compared to both T_N_ and T_CM_, while the expression of the naïve-enriched genes such as *CD38*, *TCF7*, *SELL*, *FOXP1*, *CCR7*, *NELL2*, *LTB*, *CD27* were either higher or unchanged in T_SCM_ compared to T_CM_ (Fig. 2, A and B, and data file S1). Even in a pairwise comparison the T_SCM_ compartment showed enrichment for effector-associated genes compared to both T_N_ and T_CM_ (Fig. 2, C and D, and data file S1). Further, the gene-set enrichment analysis (GSEA) using CD8-effector enriched gene set (*36*) as well as T_N_ (*vs* T_EMRA_) and T_EMRA_ (*vs* T_N_) enriched gene sets (this study), revealed that T_SCM_ is enriched for both long-term memory or naïve-associated gene signatures compared to T_CM_ and effector-specific gene signatures compared to both T_CM_ and T_N_ (Fig. 2, E and F, and data file S2). Together these results revealed that the T_SCM_ compartment shows dual signatures, where they are both stem-like as well as effector-like when compared to the other long-term memory subset, T_CM_, hence phenocopying the signatures of naïve as well as effector cells. This intriguing observation however led us to examine if T_SCM_ cells are the potential progenitors that develop into CTL lineage in the CD4^+^ T cell compartment.

**Fig. 2.**
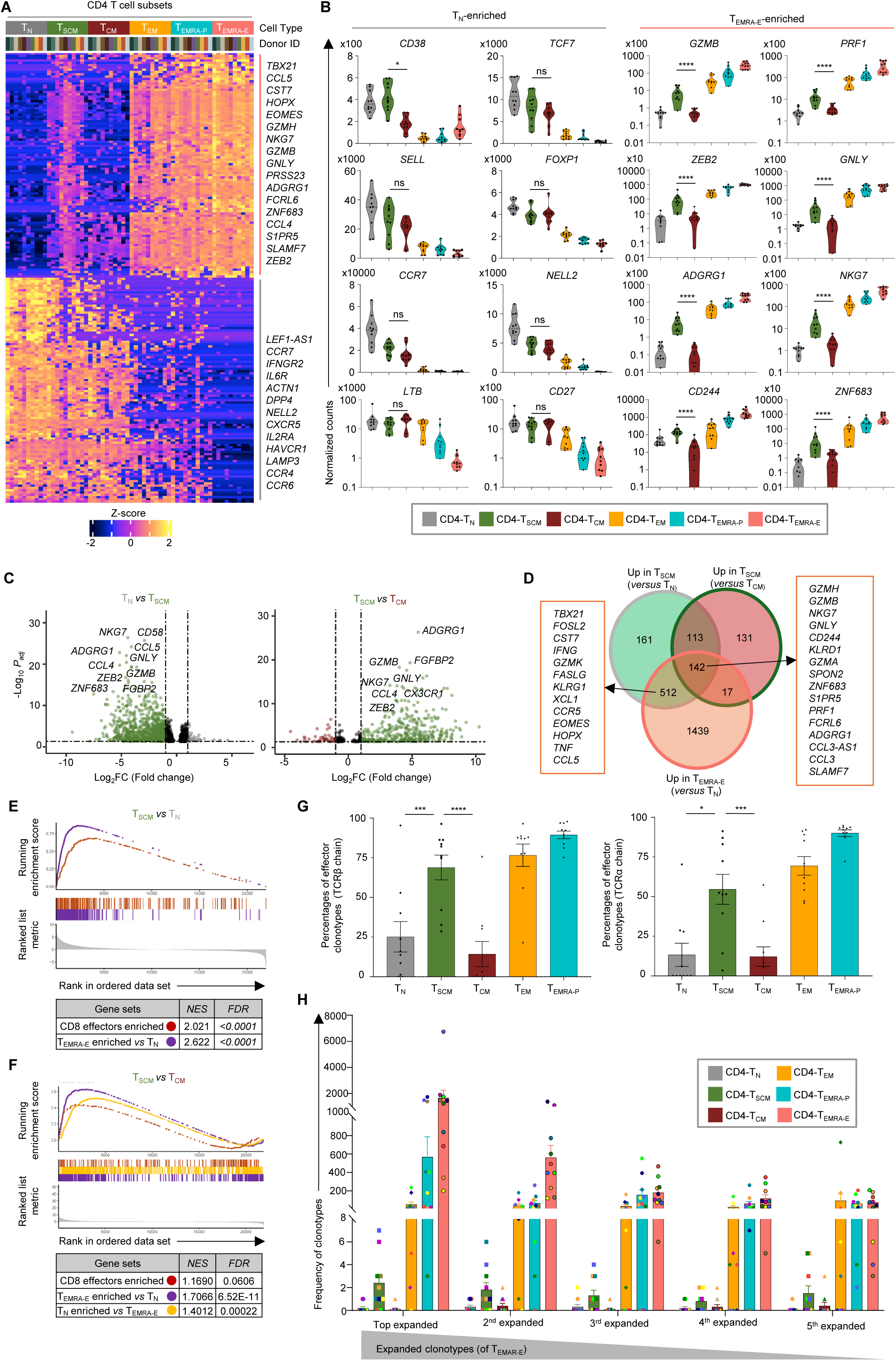
CD4^+^ T stem-cell memory (T_SCM_) shows dual signatures. (A) Heatmap of bulk transcriptomic analysis shows the row-wise z-score normalized expression of top 100 (each) significantly upregulated transcripts based on DESeq2, between T_N_ and T_EMRA-E_, visualized across the indicated CD4^+^ T cell compartments from 10 donors. (B) Violin plots of normalized counts for indicated T_N_-enriched (compared to T_EMRA-E_) and T_EMRA-E_-enriched (compared to T_N_) transcripts across the indicated CD4^+^ T cell compartments. Benjamini-Hochberg *P_adj_* *<0.05, **<0.01, ***<0.005, ****<0.001 from DESeq2 analysis. Each dot represents a donor. (C) Volcano plot for pairwise-differentially expressed transcripts based on DESeq2 between T_N_ vs T_SCM_ (left) and T_SCM_ vs T_CM_ (right), colored based on cluster where they are upregulated. (D) Venn diagram shows the transcripts enriched in T_SCM_, T_CM_ and T_EMRA-E_ subsets from the indicated pair-wise comparisons using DESeq2. Examples of overlapping genes are shown in boxes. (E-F) Combined GSEA plots show the enrichment of the indicated gene sets in T_SCM_ compared to T_N_ (E) and T_CM_ (F). Individual normalized enrichment scores (NES) and false discovery rates (FDR) are tabulated. (G) Bar graphs show the percentage the expanded clonotypes (≥3, TCRβ (left) and TCRα (right)) from T_EMRA-E_ shared by the indicated CD4 T cell subset across 10 donors. Error bars are mean ± SEM, each dot represents a donor (n=10). *P* *<0.05, *\*\** <0.01, *\*\*\**<0.005, *\*\*\*\**<0.001 from Student’s paired two-tailed T test. (H) Bar graph shows the frequency of the top 5 expanded clonotypes from the T_EMRA-E_ subset, across the CD4^+^ memory T cell subsets.

The T cells have natural barcodes in the form of TCRs, that act as their unique identifiers enabling one to trace them even when these T cells acquire different phenotypes upon an immunological event. To establish the potential lineage connection between CD4-CTLs and other CD4 memory subsets, we examined the TCR clonotype sharing between T_EMRA-E_ and other naïve and long-term memory subsets using the TCR-Seq data. Interestingly, we observed a significantly higher proportion of the expanded clonotypes of effectors (CD4-CTLs; T_EMRA-E_), were shared by the T_SCM_ compartment as compared to the other long-term memory compartment, T_CM_ and the T_N_ for both TCRβ and TCRα chains (Fig. 2G, and data file S3). Even the top 5 most expanded clonotypes of effectors were found in the T_SCM_ subset, although these clonotypes themselves were not greatly expanded in T_SCM_ (Fig. 2H, and data file S3). Over 80% and 70% of the donors share the top expanded and 2^nd^ most expanded clonotypes respectively between T_SCM_ and effectors (fig. S2C, and data file S3). Together, these results suggest a potential developmental connection between T_SCM_ and T_EMRA_ within the CD4^+^ T cell compartment.

### Identification of T_SCM-CTL_ subset within T_SCM_

We next wanted to address any potential heterogeneity in the T_SCM_ subset and identify sub-population(s) of T_SCM_ pre-committed to the CD4-CTL phenotype. To identify heterogeneity at both transcriptomic and epigenetic level, we performed scRNA-Seq and scATAC (Assay for Transposase Accessible Chromatin)-Seq assays on the T_SCM_ subset isolated from PBMCs of 6-8 donors. The scRNA-Seq analysis revealed 15 clusters, and as expected considering the stem-like nature of this subset, most of the clusters expressed long-term memory/naïve-associated transcripts such as *CD27*, *NELL2*, *TCF7* (Fig. 3, A and B, and data file S4). However, additionally these cells expressed gene signatures of other functional subsets in a cluster-specific manner (Fig. 3, A and B, and data file S4). For example, cluster 5 showed enrichment for transcripts of TFs known to be expressed by stem-cell/naïve T cells such as *ZEB1*, *BACH2*, *POU2F1*, *KLF12* etc suggesting these cells are possibly the most undifferentiated cells or show higher stemness features (Fig. 3B, and data file S4) (*41–45*). Similarly, cluster 7 showed T_REG_ specific transcripts, *FOXP3*, *IL2RA*, *TIGIT*, *IKZF2*, *CTLA-4*, *IL10RA* indicating potential stem-cell memory precursors of T_REGs_ (Fig. 3B, and data file S4). Most importantly, clusters 11 and 12 were enriched for effector-specific transcripts associated with cytolytic ability such as *GZMB*, *GNLY*, *PRF1*, and TFs *ZEB2*, *HOPX*, *ZNF683*, *TBX21*, *BHLHE40* and showed enrichment for cytotoxicity- and effector-enriched gene sets, hence representing the CTLs in the T_SCM_ subset (hereafter referred to as T_SCM-CTLs_) (Fig. 3, B to D, fig. S3A, and data files S2 and S5) (*12, 23, 46–48*). The presence of T_SCM-CTLs_ in T_SCM_ was also further validated at protein level for CTL-associated molecules such as GZMB, PRF1, GNLY, CX3CR1, CD244 and GPR56 (ADGRG1) using flow cytometry analysis (fig. S3B). Consistently, the T_SCM-CTLs_ constituted a very small fraction of T_SCM_, even in flow cytometry analysis, and were spread across the T_SCM_ subset without showing any bias for either low or high expression of CD45RA and CCR7 (fig. S3B). Although both clusters 11 and 12 showed enrichment for cytotoxicity-associated transcripts compared to other clusters, overall, the expressions were more pronounced in cluster 11 compared to 12, hinting towards probably more than one stage in the CTL development within the T_SCM_ subset (Fig. 3, B to D, and data file S4). Hence, next to understand if a subset of cells in T_SCM_ are in a poised state for the CTL lineage, and to identify the gene-expression regulation patterns of these cells, we performed scATAC-seq and identified 8 distinct clusters based on chromatin accessibility (Fig. 3E). Two clusters, cluster 6 and 7 showed overall higher mean accessibility score for peaks linked to the cytotoxicity-associated, CD4-T_EMRA_-associated and effector (*vs* naïve)-enriched gene sets, with cluster 6 showing relatively higher accessibility compared to cluster 7 (Fig. 3F, fig. S3C, and data file S2) (*12, 49*). Based on the prediction scores obtained from an integrative analysis of scRNA-Seq and scATAC-Seq data, we predicted the cells in clusters 6 and 7 to be the T_SCM-CTL_ cells identified in scRNA-seq (Fig. 3G). We then analyzed chromatin accessibility of the genomic loci that contained CTL-associated genes across the clusters using genomic tracks and calculated group gene scores that are indicative of gene activity (Fig. 3H, violin plots). Both clusters 6 and 7 showed overall higher gene activity for GZMB, CCL5, PRF1, TBX21, SLAMF7, and ADGRG1 (Fig. 3H, violin plots). Further, we found several peaks in either the gene body or in the regulatory regions, of genes such as GZMB, NKG7, CCL5, SLAMF7, TBX21 that were differentially accessible, showing a relatively higher accessibility in cluster 6 than cluster 7, and several of these peaks overlapped with histone marks associated with gene activation such as H3K4me1 and H3K27ac (highlighted area in Fig. 3H) (*50–53*). However, the chromatin accessibility of long-term memory associated genes such as CCR7, CD27, CD28 etc did not show any significant variation across these clusters (fig. S3D). Importantly, we found several peaks showing positive correlation, implicating co-accessibility, thus suggesting a probable association of these open peaks with gene expression in a co-regulated manner (Fig. 3H) (*54*). For example, peaks upstream of CCL5, SLAMF7, ADGRG1 and PRF1 in cluster 6, showed higher correlation with multiple peak/s in the gene body, indicating a possible interaction of these genomic region in the regulation of expression of their transcripts (Fig. 3H). Further, in combination with the chromatin immunoprecipitation sequencing (ChIP-Seq) data from literature, we found that several of these peaks in the cytotoxicity-related genes could be bound by the TFs of CTL relevance (EOMES, T-bet and FOSL2) (Fig. 3H) (*55–57*). We then assessed the motif enrichment across these clusters and found that the CTL-specific TFs, EOMES and TBX21 showed higher motif enrichment in cluster 6 followed by cluster 7 while the naïve/long-term memory-specific TFs, LEF1 and TCF7 showed no major noticeable difference (Fig. 3I, and fig. S3E). Interestingly, we also noted that the cluster 6 cells were predicted to have cells from both clusters 11 and 12 of scRNA-Seq data (clusters 6-11 and 6-12), indicating further heterogeneity at the level of chromatin accessibility (Fig. 3G). Genomic tracks for cytotoxicity-associated genes confirm this variability within cluster 6, where cluster 6-11 and cluster 6-12 show differential accessibility for some genes such as GZMB, SLAMF7, PRF1, while other genes such as CCL5 and TBX21 showed no variation (Fig. 3H). Together these observations suggest a tight regulation of the gene expression of cytotoxicity-related genes in T_SCM-CTL_. Overall, the scRNA-Seq and scATAC-seq suggests heterogeneity in the T_SCM_ subset and may indicate committed stem-like cells to different T cell lineages, where the T_SCM-CTLs_ are poised for the CTL lineage.

**Fig. 3.**
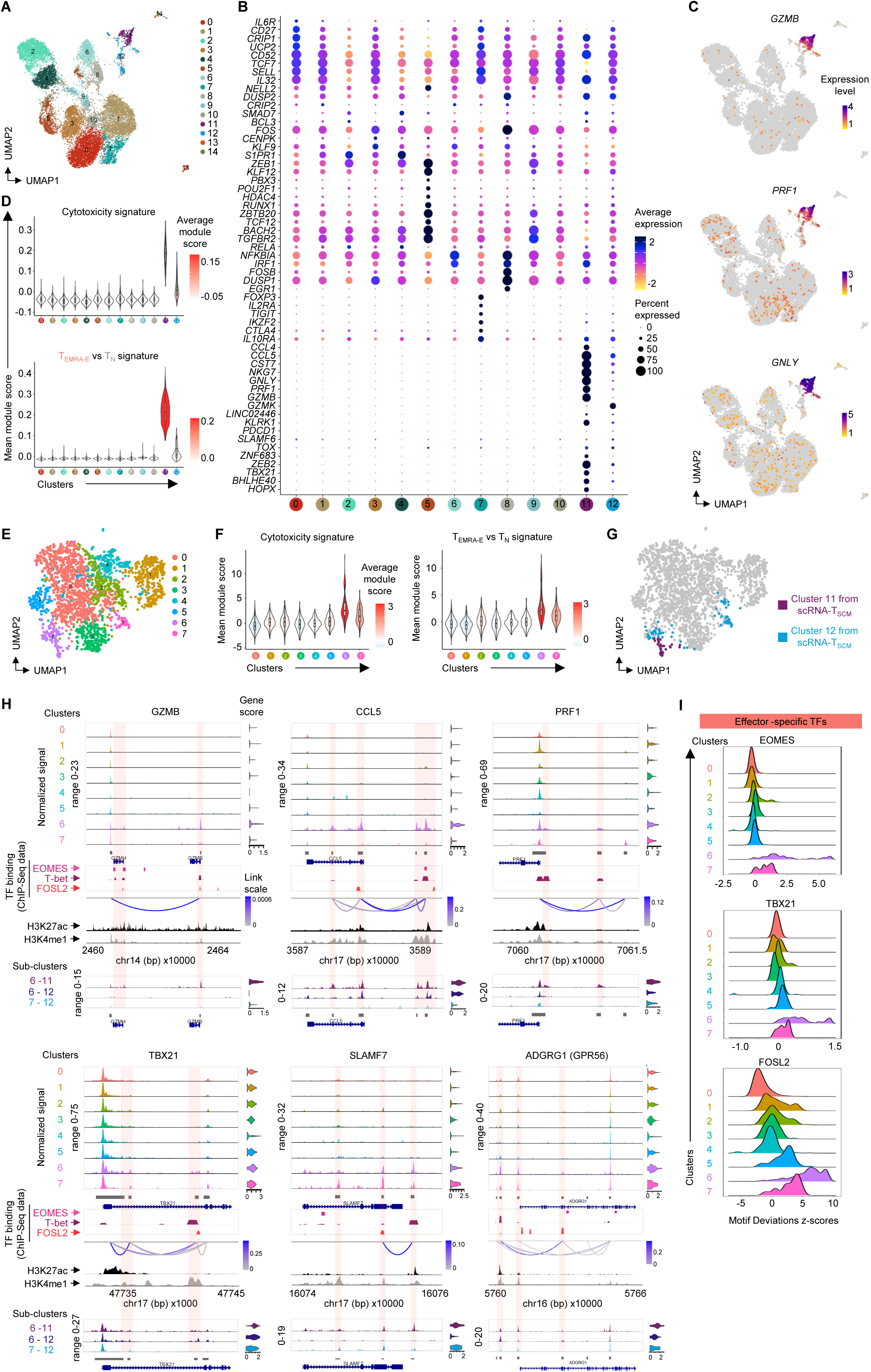
Single-cell (sc) RNA-Seq and scATAC-Seq analysis of T_SCM_ cells. (A) 2-D UMAP projection of the scRNA-Seq analysis of 17,364 T_SCM_ cells. (B) Dot plot shows mean expression (color) and percentage of expressing cell (size) of indicated differentially expressed transcripts across clusters (C) 2-D UMAP visualization of normalized expressions of indicated transcripts. (D) Violin plot for the mean signature scores (y-axis) for indicated gene modules per cluster (x-axis) painted by the average score. (E) 2-D UMAP embedding of scATAC-Seq data from 2,240 T_SCM_ cells. (F) Violin plot for the mean accessibility scores (y-axis) of the peaks linked to the signature genes for indicated gene modules per cluster (x-axis) painted by the average score. (G) 2-D UMAP embedding painted by the predicted identity of cluster 11 and 12 from scRNA-Seq analysis of T_SCM_ projected onto scATAC-seq data. (H) Chromatin accessibility of indicated gene-containing genomic loci across indicated clusters/sub-clusters, visualized using CoveragePlot (top) with group gene score as violin plots (right) and significant co-accessible peaks connected via links colored based on scores. Publicly available ChIP-Seq data for T-bet, EOMES and FOSL2 for the corresponding genomic loci shown as colored peak plots. ENCODE sourced histone modification marks of H3K27ac (black) and H3K4me1 (grey) are shown. Peaks or regions of interest are highlighted. (I) Ridge plot of chromVAR deviation (z-) scores for indicated effector-associated TFs across clusters. For (B and D), clusters with <1% of the total cells are not shown (clusters 13=167 and 14=86 cells).

Next, to identify TFs of unknown CTL function, that could play a role in early commitment to the CTL lineage, we examined the transcript expression of all the known TFs (1639) found in the T_SCM_ single-cell transcriptome data set (1163 of the 1639 known TFs in human genome) (*58, 59*). Interestingly, the hierarchical clustering based on these 1163 TFs expression identified cluster 5 closely resembling the T_SCM-CTL_ clusters 11 and 12 (fig. S3F). To identify potential TF of CTL relevance amongst them, we identified TFs that showed shared expression between clusters 5, 11 and 12, by performing DE analysis between these clusters versus the rest of the clusters (fig. S3G, and data file S4). Expectedly, the T_SCM-CTL_ clusters 11 and 12 showed relatively higher expression of known CTL-associated TFs, *ZNF683* (encoding Hobit), *ZEB2*, *EOMES*, *TBX21*, *BHLHE40*, *FOSL2* etc (fig. S3, G and H, and data file S4) (*12, 23, 46–48*). Of particular interest was a set of TFs that included *STAT4*, *NFAT5*, *REL*, *KLF12*, *CUX1*, *FOXN3*, *RUNX2*, *NFATC3* etc that were co-expressed by cluster 5 and the T_SCM-CTL_ clusters 11 and 12, with cluster 12 showing relatively higher expression than cluster 11 (fig. S3, G and H). Amongst these, some are already reported to have function in effectors; NFAT5 has recently been shown to be associated with CD8-effectors and to induce exhaustion in CD8^+^ T cells and was also one of the DEGs associated with CD4-T_EMRA-E_ subset (*vs* T_N_) in this study (data file S1) and STAT4 the known regulator of T-bet in T_H_1 lineage (*60–62*). However, many of these TFs have been shown to play role in stemness or proliferation - TCF12, a member of the basic helix-loop-helix (bHLH) E-protein, has been shown to be expressed by many cell types including T and B cells and hematopoietic stem cells and KLF12 is known to promote proliferation in cancer and Natural Killer (NK) cells (*44, 63, 64*). TCF12 is further known to form heterodimer with other BHLHE proteins, hence may form complex with BHLHE40, a TF of known CTL relevance (*47*). These observations suggest that several of these TFs may potentially regulate the CTL program, although further experimental validations are necessary to prove or disprove their role in commitment to the CTL program. Hence, cluster 5 may be the pre-T_SCM-CTL_ cluster from which the T_SCM-CTLs_ develop. Overall, the cells in clusters 5, 12, and 11 are the progenitors that chronologically appear during the development. They are likely the close parallels to the most recently described TCF1^+^PD1^+^TOX^+^ stem-like precursors of exhausted T cells and the correlates of protection described in mouse models of both chronic and acute infection (*65, 66*) (Fig. 3B).

### The CD4-CTLs are developed from T_SCM-CTLs_

We then examined if T_SCM-CTLs_ are indeed the precursors of CD4-CTL lineage by comparing the single-cell transcriptomes and scTCR repertoire of CD4-T_EMRA_ and CD4-T_SCM_ subsets from the same donors. The scRNA-seq analysis showed that most of the clusters were either T_SCM_-specific or T_EMRA_-specific, separating based on longevity and effector status (Fig. 4, A and B, and fig. S4A). The T_SCM-CTL_ clusters (Fig. 3, A to D, clusters 11 and 12) mostly mapped back to clusters 13, 14 and part of cluster 3 of the integrated data (Fig. 4C). This observation was further validated by the expression of CTL-associated (*GZMB*, *PRF1*, *GNLY*, *GPR56* and *ZNF683*) and naïve/long-term memory associated (*CCR7*, *CD27*, *CD28*, *SELL*, *LEF1*) transcripts (Fig. 4D). We then analysed the scTCR-seq data to examine the paired TCR clonotypes from these cells to establish TCR clonal connection. We noted as many as 75 unique TCR clonotypes were shared between T_EMRA_ and T_SCM_ subsets of which 56 were clonally expanded in T_EMRA_ subset (Fig. 4, E to G, and data file S5). The TCR clonotype sharing between T_SCM_ and T_EMRA_ was observed in 100% of the donors (fig. S4, B and C, and data file S5). These clonotypes themselves were not expanded in T_SCM_, with their frequency ranging from only 1 to 6, however these clonotypes were hugely expanded in the T_EMRA_ subset with few clonotype frequencies being as high as 931, 883 and 600 (Fig. 4, F and G, fig. S4C, and data file S5). Importantly, majority of these shared clonotypes were found in cluster 13 cells that carry T_SCM-CTL_ cells across all the donors (Fig. 4G, fig. S4C, and data file S5). Overall, overlaying the single-cell transcriptomic data with the paired scTCR repertoire data, we discovered a novel subset in the T_SCM_ compartment - the T_SCM-CTL_, the long-term memory subset with stemness properties, that is likely poised to differentiate to CD4-CTL lineage in an immunological event such as infection.

**Fig. 4.**
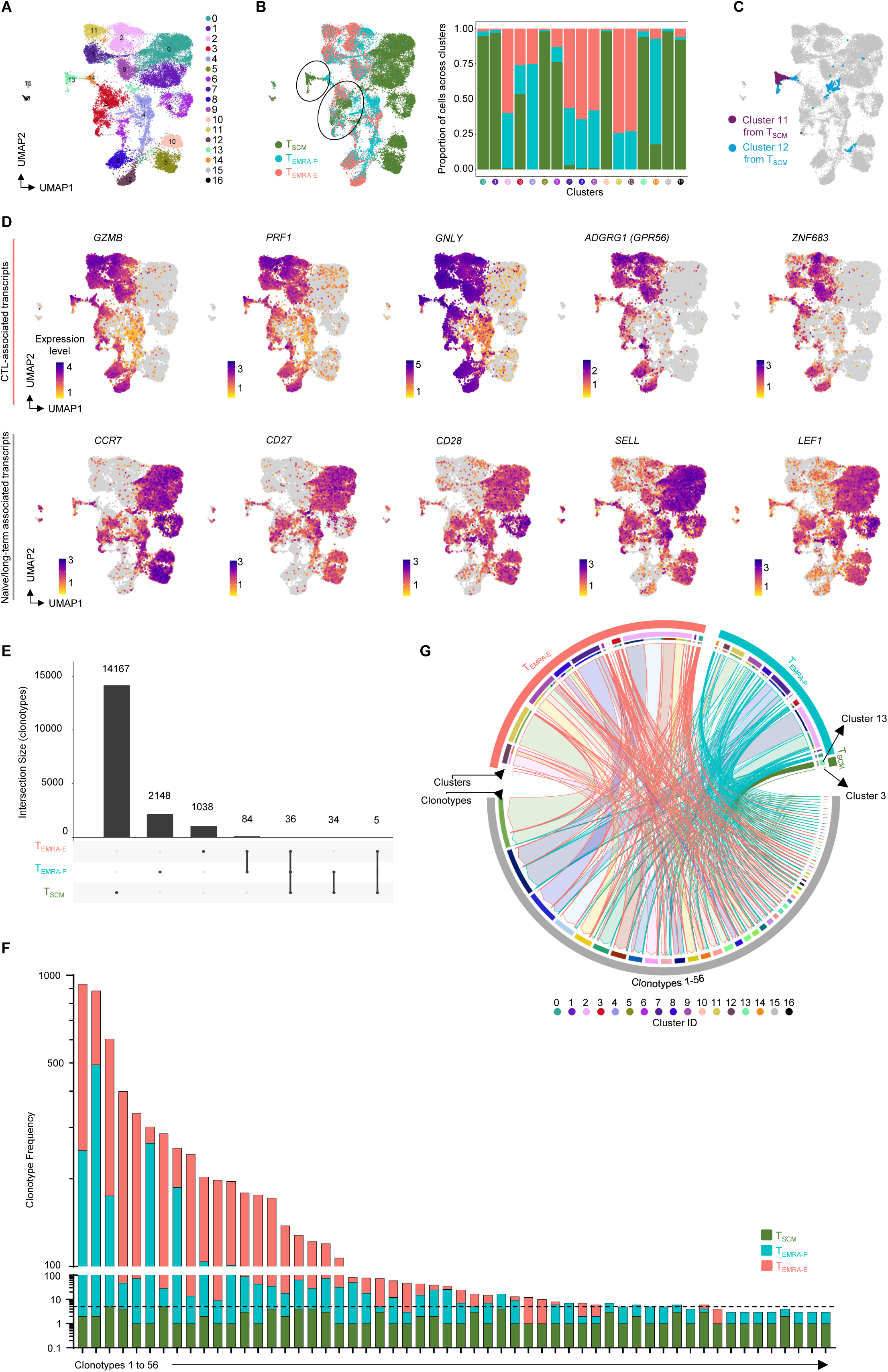
CD4-T_SCM-CTLs_ are progenitors of CD4-T_EMRA_. (A-B) Integrated 2-D UMAP embedding of scRNA-Seq analysis of T_SCM_ (17,364 cells) and T_EMRA_ (15,677 cells) shown either based on clusters (A) or origin (B-left). Stacked bar graph (B-right) shows proportion of cells from each category across different clusters (x-axis). (C) 2-D UMAP visualization of cells from clusters 11 and 12 from scRNA-Seq analysis of T_SCM_ (T_SCM-CTLs_). (D) 2-D UMAP visualization of normalized expressions of indicated transcripts from specified categories. (E) UpSet plot shows the frequency (y-axis) and distribution of unique or shared clonotypes (connected lines) across categories. (F) Stacked bar plot shows the frequency and distribution of shared clonotypes between the indicated subsets. (G) Circos plot shows distribution across clusters and categories (T_SCM_, T_EMRA-P_ and T_EMRA-E_) of 56 shared clonotypes between T_SCM_ and T_EMRA_ cells. The links between the corresponding cells with their clonotypes are colored based on the clonotype with the arrowhead pointing towards the clonotype and the border of the link colored based on the origin of the cell.

### Differentiation of CD4 naïve T cells to CD4-CTL phenotype

The molecular program that drives the naïve T_H_ cells to cytotoxic (CTL) lineage in humans is poorly understood. Hence, to systematically understand the developmental lineage of CD4-CTLs in humans, we wanted to develop an *in vitro* differentiation model to generate CD4^+^ T cells with cytotoxic potential (CD4-CTLs) from CD4 naïve T cells (T_N_) and track them over a period. Considering few overlapping gene expression patterns between CD4-CTLs and T_H_1 cells, we hypothesized that the stimulation of T_N_ cells with T_H_1-specific polarizing milieu of cytokines and blocking antibodies of other T_H_ lineages, over a period, may differentiate them to CD4-CTLs. Hence to test this hypothesis, we activated the naïve T cells via TCR stimulation using αCD3/αCD28 beads for 48 hrs and continued to culture them in a cocktail of T_H_1 or as controls, in T_H_2 and neutral polarizing cytokines and blocking antibodies along with IL2 for 8 days (2+8 days cycle) (Fig. 5A). The successful polarization of T_N_ cells to T_H_1 or T_H_2 lineage was confirmed by the expression of cytokines and TFs of their respective linages; IFNγ, T-bet and EOMES for T_H_1 and, IL4 and GATA3 for T_H_2 (fig. S5A). In alignment with our hypothesis, we observed a higher proportion of cells making cytolytic molecules such as GZMB, PRF1, and GNLY under T_H_1 compared to T_H_2 or neutral polarizing conditions (Fig. 5B). Consistent with this observation, previous reports in both human and animal models have also shown that the T_H_1 polarizing cytokines can induce the expression of GZMB and PRF1 (*67–70*). Importantly, the expression of these cytolytic markers was dynamic and was more pronounced from 2^nd^ round of stimulation (day(D)15 onwards), indicating that constant exposure to desirable cytokine milieu can polarize naïve T_H_ cells to T_H_ cells expressing cytolytic molecules (CD4-CTLs) (Fig. 5B). We then performed an integrated multi-factorial analysis that included expression of cytolytic molecules, GZMB, PRF1, GNLY and memory markers CD45RA and CCR7, different polarizing conditions, days after polarization, and donors (n=3 to 6) to understand the dynamic changes during the differentiation process using the flow cytometry data (Fig. 5, C to E, and fig. S5B). An unbiased clustering using a similar number of cells from T_H_1 and T_H_2, revealed that they were represented by distinct clusters, and the ones (7 and 12), showing relatively higher expression of the cytolytic molecules GZMB, PRF1 and GNLY, predominantly belonged to the T_H_1 group (Fig. 5, C to E, and fig. S5B). Further within the T_H_1 group, there was a progressive increase with days (D8 to D27) in the cells co-expressing cytolytic markers GZMB, PRF1, GNLY that also showed higher expression of CD45RA and lower expression of CCR7, hence suggesting that under T_H_1 polarizing condition the naïve T_H_ cells gradually acquire the expression of cytolytic molecules and appear to resemble CD4-T_EMRA_ cells (Fig. 5, D and E). The examination of other CTL markers such as CD244, KLRG1 and CX3CR1 revealed dynamicity in marker expression - while the CD244 expression gradually increased from D15 to D20 (average of 6.83% to 47.28%), the expression of KLRG1 and CX3CR1 was undetectable under T_H_1 polarized condition whereas no or very few cells expressed any of the three markers in T_H_2 or neutral polarizing conditions (Fig. 5F). Further, in an *in vitro* functional cytolysis assay the cells polarised under T_H_1 conditions showed a higher killing ability as compared to T_H_2 polarized cells (Fig. 5G). Together, these results show that naïve T_H_ cells can be differentiated to T_H_-CTL (CD4-CTL)-like cells under T_H_1 polarizing condition over a continued culture in a polarizing milieu and suggests a dynamic and gradual expression of cytolytic molecules during the development of CTL program.

**Fig. 5.**
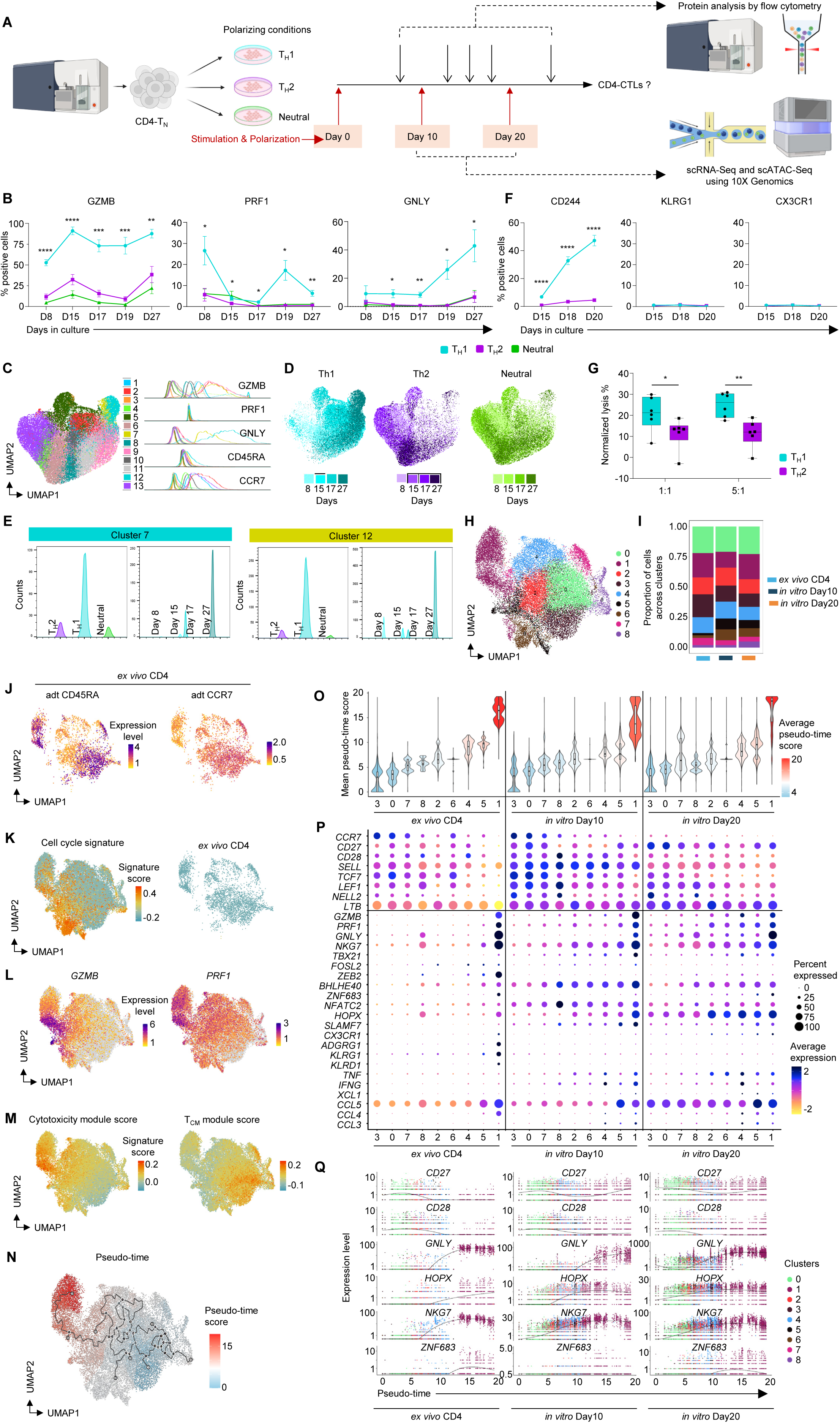
*In vitro* differentiation and polarization of naïve CD4-T (T_N_) cells to CD4-CTLs. (A) Schematic representation of experimental design for the differentiation and polarization of human naïve CD4-T (T_N_) cells into T_H_1 (n=6), T_H_2 (n=6) or neutral (n=3) conditions. (B and F) Summary flow cytometry data of indicated proteins across days comparing T_H_1-(n=6), T_H_2-(n=6) or neutral-(n=3) polarized cells. Error bars represent mean ± SEM. *P* *<0.05, *\*\**<0.01, *\*\*\**<0.005, *\*\*\*\**<0.001 from Student’s paired two-tailed T test comparing T_H_1 vs T_H_2 polarizing conditions. (C-E) Integrated 2-D UMAP visualization of multifactorial analysis using FLowJo plugin of 132,582 *in vitro* differentiated and T_H_1-, T_H_2- and neutral-polarized cells from count normalized samples across donors from days 8, 15, 17 and 27 of culture. Clusters identified using XShift (C-left) are visualized by their geometric mean fluorescence intensity (MFI) channel values for the indicated proteins (C-right). The UMAP is split by polarization and colored by days (D). Clusters 7 and 12, split based on polarization conditions or days, represented by histogram (E). (G) Box plot shows the normalized percent lysis of Raji cells (T:target cells) by T_H_1- or T_H_2-polarized and differentiated cells (E:effectors) (n=6) normalized to control Raji cells at each indicated E:T ratios. *P* *<0.05, *\*\**<0.01, *\*\*\**<0.005 from Student’s paired two-tailed T test. (H) Integrated 2-D UMAP embedding of single-cell transcriptomes of 26,815 cells from *ex vivo* CD4 T cells and *in vitro* differentiated and T_H_1 polarized T cells from day(D)10 and D20, colored by clusters. (I) Stacked bar graph shows the distribution of cells across clusters in each indicated category. (J) 2-D UMAP visualization of indicated normalized protein expression (adt) for *ex vivo* CD4 T cells. (K) 2-D UMAP projection painted by cell cycle gene signatures either together (left) or for *ex vivo* CD4 data (right). (L-N) 2-D UMAP visualization painted by normalized expressions of indicated cytotoxicity-related transcripts (L) or by indicated gene signatures (M) or by pseudo-time scores (N). (O) Violin plot for the mean pseudo-time score (y-axis) arranged based on increasing scores per cluster (x-axis) painted by the average score. (P) Dot plot shows mean expression (color) and percentage of expressing cell (size) of indicated transcripts split between clusters, grouped by origin. (Q) Scatter plot of normalized expression of indicated transcripts (y-axis) along the pseudo-time trajectory (scored along x-axis) colored based on clusters.

Since we observed a significant increase in the expression of cytolytic molecules and cytolysis of target cells under T_H_1 polarizing condition compared to T_H_2 or neutral, further to understand the overall cellular heterogeneity and the dynamic changes in the molecular program of CTL development, we performed paired single-cell RNA-Seq, and TCR-Seq on naïve cells polarized under T_H_1 condition at two different stages; D10 (potentially committed to CTL lineage) and D20 (cells with active CTL program) (Fig. 5A). To understand the developmental trajectories of these *in vitro* differentiated cells in the context of well-defined developmental memory stages, we compared them with the single-cell transcriptomes of *ex vivo* isolated unstimulated total CD4^+^ T cells from the same donors (Fig. 5H, and fig. S5C). The integrated scRNA-Seq analysis of 26,815 cells, identified 9 clusters revealing heterogeneity in differentiated cells that co-clustered with *ex vivo* CD4^+^ T cell subsets classified based on CD45RA and CCR7 as naïve (majorly cells from clusters 3 and 0, and part of 7 and 8), memory (T_CM_ + T_EM_) and T_EMRA_ (cluster 1) subsets (Fig. 5, H to J, and fig. S5, C and D). Most clusters, except clusters 5 and 6, consisted of cells from all groups, *ex vivo* CD4^+^ T cells, D10 and D20 cells, indicating the cells differentiated under the T_H_1 polarizing condition consist of cells that resemble different developmental *ex vivo* memory subsets (Fig. 5I, H and I, and Fig. 5, C and D). Interestingly, clusters 5 and 6 comprised of majorly *in vitro* differentiated cells and were enriched for cell cycle signatures, hence justifying their absence in quiescent *ex vivo* CD4 group (Fig. 5, H to J, and data file S2) (*71*). Cluster 1, that showed higher expression of CTL-associated transcripts *GZMB* and *PRF1*, included CD4-T_EMRA_ cells (CD45RA^+^CCR7^-^) from the *ex vivo* CD4 group (Fig. 5, K and L). This was further confirmed by the enrichment of signature score for cytotoxicity/T_EMRA_ in cluster 1 that showed lower enrichment for long-term memory/naïve gene sets (Fig. 5M, fig. S5E, and data file S2) (*12*). To further understand their developmental trajectories in an unbiased way, we performed pseudo-time analysis using Monocle3 (Fig. 5N) (*72*). Overlaying the pseudo-time scores, we could assign clusters to different developmental trajectories with cluster 3 & 0 being the starting point with naïve-like phenotype, and cluster 1 being the end point with T_EMRA_-like phenotype, and the rest of the clusters showing a continuity or transitioning states in memory subsets. Though cluster 1 expressed cytolytic molecules across all three groups (*ex vivo* CD4 T cells, *in vitro* differentiated cells from D10 and D20), the *in vitro* differentiated cells from other clusters also expressed cytolytic molecules, albeit at a reduced level, while the *ex vivo* CD4 group did not (Fig. 5P). Further, when we assessed genes on the pseudo-time scale, the naïve/long-term memory-associated genes (*CD28*, *CD27*) and effector-associated genes (*GZMB*, *PRF1*, *GNLY*) were expressed by distinct clusters from the *ex vivo* CD4 group, while the cells from *in vitro* D10 or D20 groups co-expressed both sets of genes (Fig. 5, P and Q). The CTL-associated TFs showed dynamic expression patterns during the differentiation process; while *TBX21*, *FOSL2*, *BHLHE40,* and *NFATC2* showed relatively more expression at D10 compared to D20, especially in clusters positioned late in the pseudo-time scales (clusters 2, 6, 4, 5 and 1), *HOPX, ZEB2* and *ZNF683* showed higher expression at D20 compared to D10 (Fig. 5, P and Q, and fig. S5F). We further noted that amongst the clusters of early pseudo-time-scale (3, 0, 7 and 8), cluster 8 showed relatively higher expression of a few CTL/effector-associated transcripts such as *GNLY*, *NKG7*, *ZEB2*, *BHLHE40*, *NFATC2*, *SLAMF7*. This prompted us to examine if cluster 8 resembles that of cluster 5 of T_SCM_ (pre-T_SCM-CTL_; Fig. 3) and the development of CTLs from naïve T cells is via pre-T_SCM-CTLs_. To this end we observed that cluster 8 is enriched for pre-T_SCM-CTL_ (cluster 5 of T_SCM_, Fig. 3 and fig. S3)-associated gene set when compared across the groups (fig. S5, G and H, and data file S4). Consistent with this observation, we further noted higher expression of transcripts associated with TFs (*STAT4*, *NFAT5*, *KLF12*, *TCF12* etc) that we predicted to be potentially functioning upstream of CTL-TFs and CTL program (fig. S5I, S3, G and H). These results provide further evidence for CD4-CTL development from T_SCM-CTLs_ that are mostly developed from pre-T_SCM-CTLs_ that are poised for the CTL program.

### The T_N_ cells polarized under T_H_1 undergo gradual change to acquire cytotoxicity program

To understand the developmental trajectory of these cells, we analysed the scTCR-Seq data from the *in vitro* differentiated cells. The TCR-Seq analysis revealed sharing of clonotypes between D10 and D20, with 17 of the 25 most-expanded clonotypes being shared, with significantly higher expansion at D20 compared to D10 (Fig. 6A, and data file S5). When the top 20 shared clonotypes were analyzed from both D10 and D20 cells across the clusters, clonotypes found in clusters 1 and 4 that are positioned later in pseudo-time scale, showed further expansion at D20 from D10 (Fig. 6B, and data file S5). We then classified these cells based on the cytotoxicity signature score as low-, moderate- and high-cytotoxic cells to examine their trajectory of CTL program development (Fig. 6C, and data file S2) (*12*). We noted a gradual increase in the proportion of clonally expanded cells from low- to moderate-to high-cytotoxic cells (Fig. 6C, and data file S5). Together, the scTCR-Seq and scRNA-Seq data suggest that the high-cytotoxic cells at D20 are close representation of *ex vivo* T_EMRA_ (CD4-CTL effector) cells. Thus, to delineate the path of cytotoxicity program development, we examined the TCR clonotypes found in high-cytotoxic cells at D20 across the cells from D10 categories. Interestingly, we observed that though majority of the cells from the high-cytotoxic group at D20 are found in the high- or moderate-cytotoxicity group of D10, a few clonotypes were also found in the low-cytotoxic group in D10 (Fig. 6D, and data file S5). In a reverse analysis, we examined the fate of the low-cytotoxicity cells at D10 during D20 to understand if they develop in a linear fashion. The majority of the clonotypes observed in low-cytotoxic cells at D10 were found in the moderate-cytotoxic group, with minor amounts in the high-cytotoxic group in D20 cells, indicating a potential linear trajectory of development (Fig. 6D, and data file S5). However, we also found a significant proportion of cells (over 7%), still remaining in low-cytotoxic group even at D20, potentially suggesting unequal cell division ensuring a small fraction of cells remain less differentiated, maintaining stemness, hence mimicking the CD4-CTL memory development *ex vivo* (Fig. 6D). Together these observations suggest linear development for CTL program as well as provide evidence for unequal division where 2 daughter cells can have different phenotype due to non-equal division. Paralleling this observation, we further noted that the moderate-cytotoxic cells share gene expression patterns of both low- and high-cytotoxic cells at an intermediate level (Fig. 6E, and data file S6). The high cytotoxic cells show higher expression of CTL-associated cytokines and chemokines (*IFNG*, *CCL4*, *CCL5* etc) along with effector molecules (*GZMB*, *NKG7*, *KLRK1*, *GNLY* etc) (Fig. 6E, and data file S6).

**Fig. 6.**
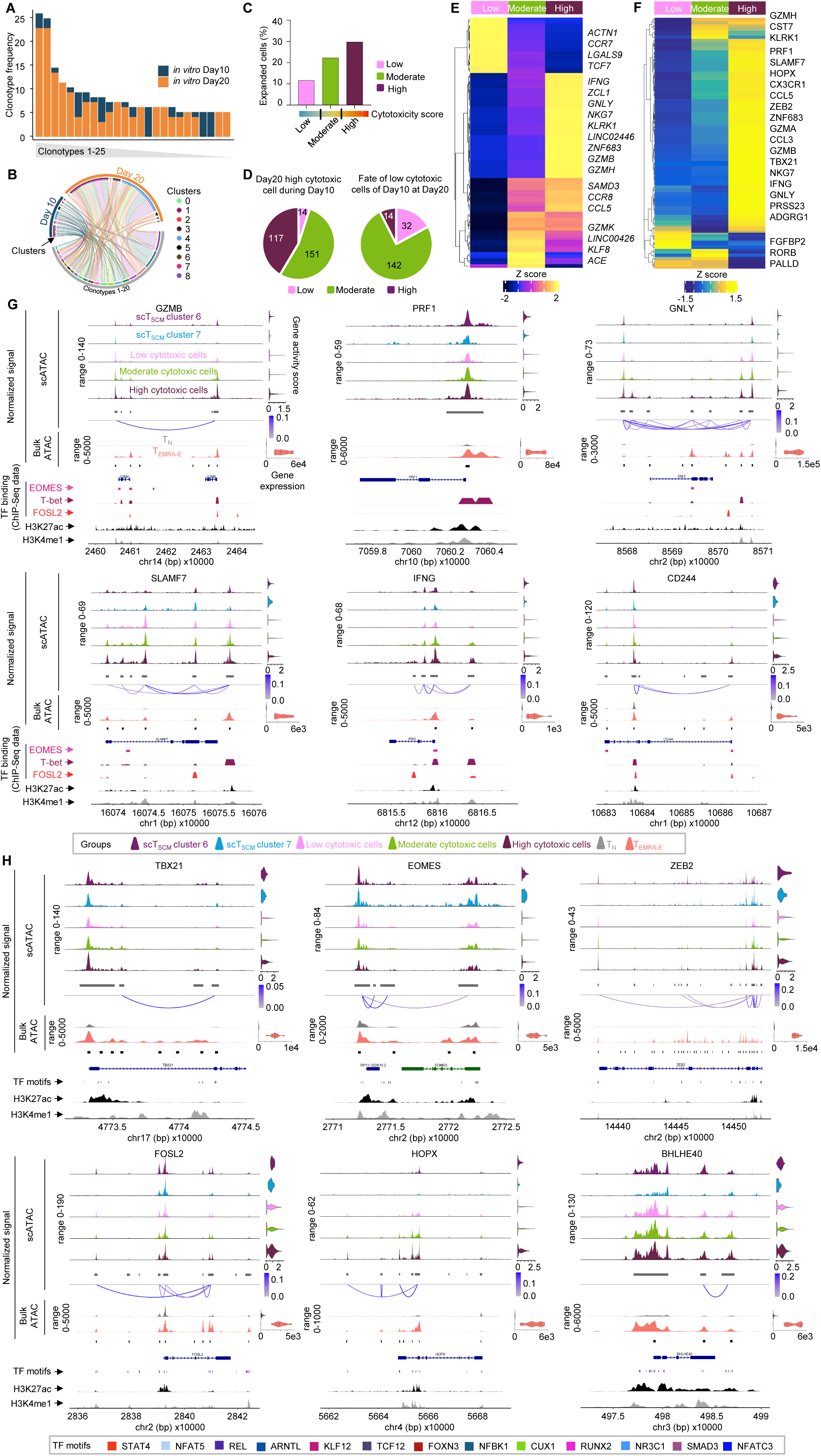
CD4-naïve T cells polarized under T_H_1 conditions undergo gradual change to acquire cytotoxicity program. (A) Stacked bar graph shows the frequency of top 25 most expanded clonotypes across days. (B) Circos plot shows top 20 shared clonotypes between D10 and D20 cells across clusters. (C) Bar graph of proportion of expanded TCR (≥2) clonotypes across cytotoxicity signature-based categorized cells. (D) Pie chart shows distribution of clonotypes grouped by cytotoxicity from the indicated time point. (E) Heatmap of scRNA-Seq analysis shows the row-wise z-score of normalized expression of the 50 most significantly enriched transcripts from each category based on the Wilcoxon Rank Sum test. (F) Heatmap of scATAC-Seq analysis shows the row-wise z-score of normalized gene activity (gene scores) of top 200 T_EMRA-E_ enriched transcripts in each category. For both (E) and (F) each column represents the average expression of all cells for a given category. (G-H) Single-cell (T_SCM_ and differentiated cells) or bulk (CD4-T_N_ and T_EMRA-E_, n=10, aggregated) ATAC-Seq analysis show the chromatin accessibility of the indicated gene-containing genomic loci, visualized using CoveragePlot with cluster-wise group gene score for single-cell open chromatin data or normalized gene expression (transcriptomic data) for bulk (T_N_ and T_EMRA-E_) data as violin plots (right) and significant co-accessible peaks connected via links colored based on scores (graded blue scale, cut-off <0.05). ENCODE sourced histone modification marks of H3K27ac and H3K4me1 are shown. Publicly available ChIP-Seq data for T-bet, EOMES and FOSL2 for the corresponding genomic loci shown as colored peak plots (G). TF of interest (different colors) having consensus motifs in peaks from scATAC data of *in vitro* differentiated cells represented as peak plot (H).

The open chromatin landscape can aid in dissecting the regulation patterns of gene expression. Hence, we performed scATAC-Seq on cells polarized under T_H_1 conditions at D10 and D20. Similar to the scRNA-Seq data, we could categorize cells as low-, moderate- and high-cytotoxicity, based on the cytotoxicity module score for peaks in scATAC-Seq data (Fig. 6F, and data file S2). The gene activity of the T_EMRA-E_-enriched genes (*vs* T_N_; data file S1), across these three categories showed that a large set of genes showed the highest activity in high-cytotoxicity group, followed by moderate- and low-cytotoxicity groups (Fig. 6F). Further, the scATAC-Seq data analysis revealed that the peaks at the regulatory regions as well as the gene body of the CTL-associated genes such as granzymes (GZMB and GZMH), perforin (PRF1), granulysin (GNLY), SLAMF7, and CD244, cytokines such as IFNG, as well as the TFs associated with CTL program TBX21, ZEB2, FOSL2, and HOPX, showed gradual opening from low-to moderate-to high-cytotoxic cells, however in few genes such as EOMES, BHLHE40 the variation between the categories was minimal especially at the TSS (Fig. 6, G and H) (*12, 23, 46–48*). These groups also show motif activity for TFs of CTL-relevance and long-term memory/naïve associated in a graded fashion, with high-cytotoxic cells showing the highest activity for TBX21, EOMES, FOSL2, while the low-cytotoxic ones showing relatively higher activity for TCF7 and LEF1 (fig. S6A). To understand the chromatin accessibility of these *in vitro* differentiated CTLs, in the context of *ex vivo* cells, we then compared them with chromatin accessibility profiles of the T_SCM-CTL_ clusters (clusters 6 and 7; Fig. 3, E to G), as well as the CD4-T_N_ cells from which the programming to the CTL lineage was initiated, and the CD4-T_EMRA-E_ cells, the bona fide CD4-CTL effector cells, and further, overlayed it with the histone activation marks such as H3K4me1 and H3K27ac from the ENCODE database (Fig. 6, G and H) (*50–53*). The open peaks at the genomic loci of the CTL-associated genes in *in vitro* differentiated cells matched with that of CD4-T_EMRA-E_ subset and were mostly missing from the CD4-T_N_ subset, suggesting the naïve cells that were differentiated to CD4-CTLs *in vitro* have CTL program activated (Fig. 6, G and H). For most of the genes, the open chromatin patterns of the moderate-cytotoxic group matched with that of T_SCM-CTL_ cluster 6 and low-cytotoxic group matched with that of cluster 7, although we noted few peak-specific variations; EOMES showed no major difference between categories and for HOPX, the T_SCM-CTL_ cluster 6 resemble low cytotoxicity (Fig. 6, G and H). Further, the peaks at the TSS of CTL-associated genes GZMB, PRF1, SLAMF7, IFNG and CTL-TFs TBX21, EOMES, ZEB2, FOSL2, BHLHE40, correspond to overall activation marks H3K27ac, and H3K4me1, an activation mark associated with promoters and enhancers (Fig. 6, G and H) (*51–53*). We then compared the binding of CTL-TFs such as T-bet, EOMES and FOSL2 in these peaks using ChIP-Seq data from literature and found that many of these chromatin regions can be bound by one or more of these TFs (T-bet, EOMES and FOSL2) (Fig. 6G) (*55–57*). Several peaks in the intronic regions or the upstream of TSS of the CTL-associated genes and TFs, corresponded to H3K4me1 and/or H3K27ac, indicating that these peaks may be probable enhancers. Considering several of these peaks can further, potentially be bound by CTL-TFs, T-bet, EOMES and FOSL2 (eg. SLAMF7, IFNG, GNLY), strengthens this observation (Fig. 6G). The co-accessibility analysis (cicero-generated links) of these peaks revealed no specific pattern based on the position of these peaks with respect to the gene body (upstream, downstream or intronic regions) for the co-accessibility, indicating a complex and distinct regulatory mechanism employed by these cells in activating different genes in the CTL program (Fig. 6, G and H). Furthermore, comparing the intensity of peaks between low-cytotoxic cells and cells from T_SCM_ suggests that the *in vitro* differentiated cells indeed go through a phase where they represent the T_SCM-CTLs_.

We next examined if the TFs that we identified in the scRNA-Seq data from T_SCM_ (clusters 5, 11 and 12) and *in vitro* differentiation (cluster 8), could be regulating the CTL program probably through functioning upstream in the signalling cascade of CTL-TFs, by examining the potential binding of these TFs in regulatory regions of CTL genes. Through scanning of the peaks of the CTL-associated genes, we examined if these genes contain a binding site for any of the 13 TFs (of the 29, where consensus motif was available (STAT4, KLF12, TCF12, RUNX2 etc)) in the regulatory regions, especially the promoters (fig. S3H, and S6B) (*73*). Several CTL-associated genes contained the binding sites for one or more of the 13 TFs in their promoter regions and interestingly, several of them had binding sites for more than one TF (Fig. 6H, and fig. S6, B and C). To specifically examine if these can regulate the CTL-TFs such as TBX21, EOMES, FOSL2, HOPX, BHLHE40, ZEB2, which may in turn regulate the CTL program, we looked for consensus motif for binding of these 13 TFs in the genomic region of the CTL-TFs. We observed that the genomic region around the CTL-TFs had motifs for several of these TFs upstream and across the gene body including intronic regions overlapping with H3K4me1, probable enhancers (Fig. 6H). STAT4 has been shown to regulate T-bet and turn on the T_H_1 program (*61, 62*). As expected, peaks in the genomic region of TBX21, which are selectively open in T_EMRA-E_, cluster 6 of T_SCM_, and *in vitro* differentiated cells, contained STAT4 binding sites (Fig. 6H). Further, the peaks in the gene body of TBX21, additionally contain binding sites for KLF12 and TCF12, and these peaks show a positive correlation among them. Similarly, the TSS of EOMES has binding sites for TCF12 and NFATC3, while upstream peaks have binding sites for REL, STAT4 and KLF12, suggesting potential regulatory roles of these TFs for EOMES activity. Consistently, gene body or, upstream or downstream regulatory regions for other CTL-TFs like ZEB2, FOSL2, BHLHE40, HOPX and ZNF683 have several peaks with binding sites for multiple of these 13 TFs, with varying degree of openness across *in vitro* differentiated cells, *ex vivo* T_SCM-CTLs_ or T_EMRA-E_ (Fig. 6H). These peaks mostly show co-accessibilities amongst them and show a graded pattern of openness when compared across developmental stages and overlap with H3K27ac or H3K4me1 marks (Fig. 6H). Together these observations suggest a possible regulation of the CTL-TFs via different TFs binding to different regulatory regions based on the developmental stage under consideration, and thus may regulate the overall CTL program either by directly regulating CTL-associated genes or the CTL-TFs (fig. S6C). However, these observations need further detailed validation using ChIP-Seq.

## DISCUSSION

The CD4-CTLs though have been identified in many diseases such as infectious diseases, a variety of cancers, auto-immune disorders etc, in both human and animal models, their functional relevance has not been very well established mainly due to their rarity in the periphery and inaccessibility due to tissue residency in humans. Considering their protective nature against many infectious diseases, their proportions can be a valuable read out of correlates of protection in natural infection as well as upon vaccination, hence, can be utilized to test the vaccine efficacy against these diseases. However, although CD4-CTLs have been shown to express many CTL-associated genes, they have not been studied in parallel to the classically known CTLs, the CD8-CTLs, hence their functional relevance has remained questionable. By studying both of these CTLs parallelly from the same donors in similar settings, we have clearly shown that they are indistinguishable from each other with respective to their transcriptomic profile and TCR clonal expansion, as well as cytotoxicity-associated protein expression (GZMB, PRF1, GNLY, CX3CR1, GPR56 etc) was at par (Fig. 1). Hence, these results suggest that an exploration of both CD4-CTLs and CD8-CTLs’s induction during vaccination and cell-based therapies may provide better outcomes. These studies in fact, provide hope for exploring the CD4-CTLs for therapeutic interventions, where the immune evasion by the pathogen happens to avoid CD8 T cell mediated killing.

The lack of understanding of the developmental lineage of CD4-CTLs, unlike CD8-CTLs, has been a major hurdle in exploring their therapeutic value (*74–78*). Hence in this study our approach was to first identify a long-term memory subset that can develop into CD4-CTL effectors and memory in response to any immunological insult. Through an integrative analysis of multi-omics approaches we identified an extremely small subset within a rare memory compartment with stemness properties (T_SCM_) that is poised for CD4-CTL lineage, the T_SCM-CTL_ subset (Fig. 2 and 3). The TCR clonal sharing provided evidence for the development of CD4-CTLs from T_SCM-CTL_ (Fig. 4) (*12*). The dynamic nature of CTL-associated TFs may indicate that different stages of CTL program can be regulated by different sets of TFs and hence, they may modulate the expression of distinct groups of genes in the CTL program, with TFs such as Hobit and HOPX regulating the terminal differentiation of CTL-effectors (Fig. 5, P and Q) (*12, 23, 29, 46*). Further, a deep dive into the heterogenous T_SCM_ subset revealed a separate trajectory for CTL development with cells at different stages of development (T_SCM-CTLs_ and pre-T_SCM-CTL_ clusters; 5->12->11). These cells appear to be the close human CD4^+^ T cell counterparts of the most recently described precursors of CD8^+^ T cells that adapt to both chronic and acute infections and are the precursor of exhausted CD8^+^ T cells (*65, 66*). Though not in the context of CD4-CTLs, but similar cells have been described in CD4^+^ T cells in transplantation and cancer immunity in mouse models (*79, 80*). Thus, establishing the potentially important role played by these stem-like cells in health and disease. Further, we identified a set of 29 TFs expressed by preT_SCM-CTLs_ and the effectors that may function upstream of the cytotoxicity program. A well-known TF, STAT4 that has been shown to function upstream in the signalling cascade of CTL-associated TF, T-bet, being part of this group of 29 TFs, further strengthens their potential role in the CTL program’s initiation and/or development (*61, 62*). However, knock-down or over-expression studies along with assays that can establish occupancy of these TFs on the chromatin may provide confirmatory evidence for their role in the CTL program of both CD4^+^ and CD8^+^ T-cells.

One of the major roadblocks in exploring CD4-CTLs in clinical utilization has been their much smaller number compared to CD8-CTLs in the periphery, despite them being equally potent killers (*12, 13*). This and other studies have demonstrated that naïve CD4 T cells can be differentiated and polarized *in vitro* to CD4^+^ T cells expressing CTL-associated genes such as GZMB, PRF1, GNLY etc (*70*). Furthermore, using single-cell multi-omics we show that one can generate CD4^+^ T cells with cytolytic potential *in vitro* that are at different developmental stages. In fact, different categories within the *in vitro* differentiated cells mimicked different *ex vivo* subsets; while some mimicked the T_EMRA-E_, others, either mimicked earlier or later stage of the stem-like memory pre-committed to the CTL lineage, thus, representing variety of stages of CTL development. Most interestingly, the *in vitro* polarized CD4-CTLs while maintaining the cytolytic ability, showed signatures of long-lived memory phenotype, thus overcoming one of the major challenges with *ex vivo* isolated effectors that are unable to proliferate beyond a point due to anergy or exhaustion (*81, 82*). Thus, the system developed here can be further explored for therapeutic applications. The generation of a long-lived memory subset, that has the potential to quickly differentiate to effectors has always been considered an important goal in both natural infections and vaccinations to maintain long-term immunity against any pathogen, so that in the event of reinfection from the same pathogen, the long-term memory can quickly differentiate to effectors and provide protection against the invading pathogen. Overall, in this study we show the potential of CD4-CTL effectors in direct comparison with CD8-CTL effectors and, also delineated the developmental lineage of CD4-CTL from naïve to T_SCM-CTL_, a stem-like CD4-CTL memory subset, to CD4-CTL effectors. The data and the knowledge base generated in this study will pave the way for exploring the therapeutic potential of CD4-CTL effectors in a variety of diseases including infectious diseases, autoimmune disorders and cancers.

## MATERIALS AND METHODS

### Study subjects

Buffy coat samples were obtained from healthy donors who donated blood in Safdurjang Blood Bank during the period from October 2019 to June 2024. The donors consented to the use of the plasma and buffy coat samples for research. Donors were HIV-negative and had no history of Hepatitis C infection. The median age was 29 years (ranged from 19 to 50 years), and all these donors were male. Approval for the use of this material for research was obtained from the institutional human ethics committees from both the National Institute of Immunology, New Delhi and Vardhman Mahavir Medical College (VMMC) and Safdarjung Hospital, New Delhi, India. Paired plasma samples obtained from the blood bank and were stored in -80°C until further use.

### PBMC isolation

Peripheral blood mononuclear cells (PBMCs) were isolated from the buffy coats using density gradient centrifugation using Ficoll-Paque Premium (GE Healthcare Biosciences). PBMCs were cryopreserved in 90% fetal bovine serum (FBS) supplemented with 10% di-methyl sulfoxide (DMSO).

### Flow cytometry

Cryopreserved PBMCs were thawed, blocked using human IgG (1:25) and stained using fluorescence conjugated cell surface protein cocktails in MACS buffer (PBS supplemented with 2% FBS and 2 mM EDTA) and were analysed for protein expression using flow cytometry. For intracellular protein staining, surface-stained cells were washed and fixed using Cyto-Fast^TM^ Fix/Perm Buffer Set (cytokines and other intracellular proteins) or True-Nuclear^TM^ Transcription Factor Buffer Set (for TFs) from BioLegend, followed by staining with cocktail of antibodies. The stained cells were analysed using BD LSR Fortessa or BD Symphony A5. For studies involving degranulation and cytokine secretion assays, cryopreserved PBMCs were thawed and rested for 6 hrs at 37°C with 5% CO_2_ before stimulation using CytoStim (Miltenyi Biotec) for 6 hrs. The degranulation marker LAMP1 (CD107a) antibody was added during the stimulation and Brefeldin A was added after 2 hrs of stimulation to stop vesicular export. After 6 hrs of stimulation, the cells were processed for analysis using flow cytometry. All flow cytometry data were analysed using BD FlowJo v10.8.1, and geometric-mean fluorescence intensity along with population percentages were exported and visualized using GraphPad Prism v10.2.2.

T cell populations of interest were sorted from PBMCs after surface staining, using BD FACS Aria Fusion in collection media (1:1 FBS:PBS). For cell culture, the cells were washed and cultured as per the experimental conditions. For downstream genomics experiments, the cells were collected in collection media supplemented with 1:100 RRI (Recombinant RNase Inhibitor, Takara). For experiments involving bulk RNA, the sorted cells were washed with 1X PBS before lysing using TRIzol reagent (Invitrogen) and stored at -80°C for RNA extraction later.

### Bulk RNA-Seq

The cryopreserved TRIzol samples were thawed and used for RNA isolation using RNeasy Micro kit (Qiagen) as per manufacturer’s recommendations. Bulk RNA-seq was performed as described previously using Smart-seq2 method (*12, 83*) with 3 ng of total RNA. Briefly, the RNA was reverse transcribed, and cDNA was pre-amplified for 15 cycles followed by size selection using 0.8X AmpureXP beads (Beckmann Coulter). Quality control was performed using Fragment Analyzer 5200 (Agilent) and quantified using Quant-iT^TM^ PicoGreen^TM^ dsDNA Assay kit (Invitrogen). Final libraries were prepared using Nextera XT DNA Library Prep kit (Illumina, #FC-131-1096) following manufacturer’s protocol using 1.5 ng of pre-amplified cDNA. Libraries were then pooled and sequenced on NovaSeq 6000 (Illumina) for 150-bp paired-end sequencing.

### Bulk RNA-Seq data analysis

Raw demultiplexed FASTQ files (n=70; 10 donors, 7 cell types) were adaptor trimmed using trim_galore (https://github.com/FelixKrueger/TrimGalore) and mapped to GRCh38 (hg38) human reference genome using STAR. QC was performed using fastqc and count files were generated using HTSeq (*84*). DESeq2 was applied on depth normalized gene counts obtained from HTSeq using design = “∼0 + condition + donor” (*85*). Genes with counts < 100 across all samples taken together, were removed from downstream analysis. Pairwise cell type comparisons were performed, and genes were considered differentially expressed if it satisfied the parameters Benjamini-Hochberg adjusted *P* < 0.05 and ≥2-fold change. Annotations for gene biotype were imported from EnsDb.Hsapiens.v86.

*Visualization*: PCA (Principal Component Analysis) plots were generated on most variable genes from variance-stabilization-transformation (*vst*) applied data (Fig. 1), or, based on CD4-T_EMRA_ enriched (vs T_N_) gene list (Fig. 2, data file S1). Heatmaps were generated using ComplexHeatmap package with cell type and donor information added as part of top_annotation to maintain uniformity across heatmaps (*86*). EnhancedVolcano (https://github.com/kevinblighe/EnhancedVolcano) package was used for generating volcano plots of differentially expressed genes between cell types. Violin plots to represent normalized counts across cell types were generated using GraphPad Prism v10.2.2.

### Gene set enrichment analysis (GSEA)

GSEA was used to assess whether specific gene signatures were significantly enriched between two groups, as previously described (*35*). ClusterProfiler was used to perform GSEA between two cell types using ranked genes from DESeq2 comparisons (data file S1) and gseaplot2 from enrichplot package was used to visualize the GSEA plot (*87, 88*). The gene sets from literature used for GSEA are mentioned in data file S2.

### Bulk TCR repertoire sequencing (TCR-Seq)

Bulk TCR-Seq was performed as described previously (*12, 39*). Briefly, 5 ng of total RNA was reverse transcribed using SMARTScribe Reverse Transcriptase (Takara) with TCRα and TCRβ gene specific primers followed by PCR amplification of 20 cycles using Q5 Polymerase (NEB). TCRα and TCRβ specific libraries were prepared from the reaction independently by a 2^nd^ PCR amplification using NEBNext Ultra II DNA Library Prep Kit for Illumina following manufacturer’s recommendations. Pooled libraries were sequenced using NovaSeq 6000 (Illumina) for 150-bp paired-end sequencing.

### Bulk TCR-Seq data analysis

Raw FASTQ files were demultiplexed, mapped, and analysed using MIGEC software with default settings (*38, 39*). For clonotype analysis, the V segment, followed by the entire CDR3 region (nt), and the J segment was considered for TCRα, while for TCRβ, additionally the D segment was also considered, with the counts taken from the output of FilterCdrBlastResults of the MIGEC pipeline (*39*). caldiversitystats from VDJtools package was used for calculating the Shannon-Weiner diversity index, while calcsegmentusage was used to calculate the usage of V and J segments across samples (*38*). A clonotype was considered expanded if the frequency of occurrence was ≥ 3 in the individual sample (data file S3).

### Bulk assay for Transposase-Accessible Chromatin using Sequencing (ATAC-Seq)

Largely, Omni ATAC-Seq protocol was followed with slight modifications (*89*). Cells sorted in collection media (1:1 FBS:PBS) were counted, and 55,000 cells were taken in LoBind 0.6 mL microcentrifuge tubes (Eppendorf) and washed with 0.5 mL chilled PBS. Cell pellet was resuspended in 50 uL chilled freshly-made lysis buffer (10 mM Tris-HCl pH 7.4, 10 mM NaCl, 3 mM MgCl_2_, 0.1% IGEPAL-CA630, 0.1% Tween-20, 0.01% Digitonin, 1X Protease Inhibitor, 20 mM NaBu) supplemented with 1 U/uL recombinant RNase Inhibitor (RRI, Takara). After 5 minutes of lysis on ice, 0.2 mL of wash buffer (10 mM Tris-HCl pH 7.4, 10 mM NaCl, 3 mM MgCl_2_, 0.1% Tween-20) supplemented with 1 U/uL RRI, was added and centrifuged. Pelleted nuclei were then transposed using Tn5 transposase from Illumina, in a 50 uL reaction mix supplemented with 1X PBS, 0.1% Tween-20 and 0.01% Digitonin, at 37°C for 30 minutes. Tagmented DNA was purified using Zymo DNA Clean and Concentrator-5 kit followed by amplification using barcoded Nextera XT primers with KAPA Hi-Fi HotStart Ready Mix. After the initial 5 cycles of amplification, cycle threshold determination (CTD) was performed using SYBR and ROX. Based on the CTD, additional cycles (CTD-2 cycle) were performed on the amplified DNA. Final libraries were purified and size-selected using double sided purification by Ampure XP beads (0.5X / 1.8X). Quality control and quantification was performed using Fragment Analyzer 5200 (Agilent) and Quant-iT^TM^ PicoGreen^TM^ dsDNA Assay kit (Invitrogen) respectively. Pooled libraries were sequenced using NovaSeq 6000 (Illumina) for 150-bp paired-end sequencing.

### Bulk ATAC-Seq data analysis

Raw demultiplexed FASTQ files (n = 20; 10 donors, 2 cell types – CD4-Naïve (T_N_) and CD4-T_EMRA-E_) were quality filtered, and adaptor trimmed using trim_galore (https://github.com/FelixKrueger/TrimGalore), and trimmed reads were mapped to the GRCh38 (hg38) human reference sequence using Bowtie2 (*90*). Publicly available pipelines from snakemake (https://github.com/epigen/atacseq_pipeline) and nf-core were followed to analyse ATAC-Seq data. De-duplicated reads (using picard (http://broadinstitute.github.io/picard) markduplicates) were filtered for reads mapping to ENCODE blacklisted region, mitochondrial genome and low-quality reads (bedtools and samtools) (*91, 92*). Reads passing all quality checks were called for peaks using MACS2 narrowPeak caller (*93*). BAM files for each cell type across 10 donors were merged using samtools. Merged BAM files were converted to BigWig files (using UCSC’s bedGraphToBigWig) for visualization using genome browser. The BigWig files were imported into Signac and visualized using CoveragePlot (*49*).

### *In vitro* differentiation of naïve CD4 T cells

*Ex vivo* isolated CD4^+^ naïve T cells (T_N_) from 3 to 6 donors were activated using αCD3/αCD28 DynaBeads Human T-Activator (Invitrogen, #11131D) for 48 hrs and cultured for 8 more days in T cell media (IMDM supplemented with 5% fetal bovine serum (FBS), 2% Human Serum) along with 40 U/mL of IL-2 cytokine (BioLegend) and 50 µg/mL of Gentamicin antibiotic (Invitrogen) along with either T_H_1 (n=6 donors), T_H_2 (n=6 donors) or neutral (n=3 donors) polarizing conditions; T_H_1 polarizing condition: 5 ng/mL IL-12 cytokine (R&D System), 10 ng/mL IFN-γ cytokine (BD Biosciences) with 10 µg/mL αIL-17 mAb (eBio64CAP17; eBiosciences) and 5 µg/mL αIL-4 mAb (clone 3007; R&D System) and, T_H_2 polarizing condition: 10 ng/mL IL-4 cytokine (R&D System) with 10 µg/mL αIL-12 mAb (clone 24910; R&D System), 10 µg/mL αIL-17 mAb (eBio64CAP17; eBiosciences) and 10 µg/mL αIFN-γ mAb (clone NIB42; BD Biosciences). Cells were restimulated with αCD3/αCD28 DynaBeads when they stopped to proliferate (10 days: 2 days stimulation + 8 days culture). A total of 3 stimulation cycles were performed and the culture was continued for a total of 30 days. Flow cytometry analysis for protein expression was carried out on indicated days. To validate polarization to the desired phenotype, cytokine (IFNγ and IL4) and TF profiling (T-bet, EOMES and GATA3) was performed. For cytokine analysis, cells were activated using PMA/Ionomycin for a total of 4 hrs with addition of Brefeldin A at 2 hrs to stop vesicular export, followed by flow cytometry analysis. Single-cell transcriptomic (scRNA) and scATAC-seq was performed on T_H_1 polarized cells on day 10 (D10) and day 20 (D20).

Integrated multi-factorial analysis of flow cytometry data across multiple days were analysed for co-expression after concatenating all samples across days and polarizing conditions, post down-sampling to exactly same number of live- and singlet-gated cells across samples using BD FlowJo software (v10.8.1). The concatenated file was subjected to uniform manifold approximation and projection (UMAP) reduction (arXiv:1802.03426v3) of multi-parameter protein data using default settings in FlowJo and a KNN density estimation-based clustering algorithm was performed using XShift with default settings to identify clusters based on protein co-expression (*94*). Z-score expression was mean centered across channel values and graphs were exported from FlowJo.

#### Killing assay

The target cell killing potential of the *in vitro* differentiated cells was examined by co-culturing the T_H_1 or T_H_2 polarized cells from day 18 (D18) with the B lymphoblastic cell line Raji for 6 hrs in a round-bottom 96-well plate in Effector:Target (E:T) ratios of 1:1 and 5:1. The T cells were activated and engaged with the B cells using CytoStim (Miltenyi Biotec) that acts as superantigen. After 6 hrs of co-culture, the cells were stained with fluorescent labelled CD19, CD3 and Annexin V (AV) with Propidium Iodide (PI). The Raji cells and T cells were demarcated by the surface expression of CD19 and CD3 proteins respectively, and the apoptotic Raji cells were identified as CD19^+^AV^+^ cells. The percentage of lysis was calculated by normalising to the control Raji cells incubated in a similar condition without the addition of T cells.

### Single-cell transcriptomic and single-cell TCR-seq analysis

Single cell transcriptomic and TCR repertoire profiling was carried out using 10X Genomics 5ʹ Immune Profiling solutions. Briefly, depending on the experimental requirements, either cryopreserved PBMCs were thawed for *ex vivo* experiments, or *in vitro* differentiated cells were taken from culture and stained with hashtag oligo-conjugated antibodies (BioLegend) to assign donor identities and cells of interest were sorted based on experimental requirement after staining with fluorescence conjugated cell-surface protein antibodies. *Ex vivo* sorted or in vitro cultured cells were thoroughly washed using 1X PBS and maintained at a viability >95%. To analyse a limited repertoire of cell surface proteins, the cells were further stained with oligo-tagged cell surface protein antibodies (TotalSeq C antibodies, BioLegend) based on experimental requirement. For scRNA-Seq of CD4-T_EMRA_ (Fig. 4), T_EMRA-P_ and T_EMRA-E_ were sorted individually, followed by staining of T_EMRA-E_ with oligo-conjugated αCD4 TotalSeq C antibody to be able to demarcate them in downstream analyses, and pooled in 1:1 ratio. About 25,000 cells per lane were loaded onto Chromium NextGEM Chip G or Chip K depending on kit version v1.1 or v2 respectively. Manufacturers’ recommendations were followed to generate libraries of captured cells for transcriptome, TCR repertoire and cell surface proteins. Libraries were sequenced on NovaSeq 6000 or NextSeq 2000 (Illumina) as per the manufacturer’s recommendations based on the kit version or 150-bp paired-end sequencing.

### Single cell RNA-Seq analysis

Demultiplexed raw FASTQ files corresponding to gene expression, associated V(D)J and cell surface protein libraries from each lane of 10X Genomics run were mapped to the human reference genome GRCh38 using 10X Genomics’ cellranger (v7.1.0) multi pipeline. Individual 10X Genomics run was processed independently using the R-based Seurat package (v5.0.1) before integrating multiple datasets to mitigate batch effects or were merged if the experiments were performed concomitantly (*95*). Cells from different donors were demultiplexed based on normalized hashtag oligo (HTO) count matrix using MULTIseqDemux with default parameters, and the same was used to detect and remove doublets from the datasets (*96*). Quality control parameters were determined based on the distribution of gene count and UMI (unique molecular identifier) counts for each dataset and low quality cells were filtered out: *ex vivo* CD4 data – gene count and UMI count <500; *in vitro* differentiated cells data – gene count <1500 >8000 (D10), <500 >7000 (D20) and UMI count <3500 >45000 (D10), <1000 >40000 (D20); *ex vivo* T_SCM_ and T_EMRA_ data – gene count <500 >3500 and UMI count <500 >17500. Cells were further filtered out based on percentage of mitochondrial reads: virus antigen-activated cells <15%, *in vitro* differentiated cells <20% and, *ex vivo* sorted T cells <10%. Post quality control, individual datasets were normalized using SCTransform with vst.flavor “v2” along with regressing out percentage of counts belonging to mitochondrial genes or ribosomal genes (*97*). While analysing a relatively homogeneous T cell population, to avoid the dominant effect of T cell receptor (TCR) genes for clustering analysis, all *TR(A/B/D/G)(V/J/C)* genes were removed from the transcriptome dataset prior to calculating principal components (PCs). A shared nearest neighbor (SNN) graph was constructed using FindNeighbors function using the most significant PCs determined empirically using ElbowPlot, and then clusters were determined using FindClusters function using the SLM algorithm. Cluster resolution was chosen empirically across datasets using clustree function on the principle that each cell cluster should express a unique group of genes. A uniform manifold approximation and projection (UMAP) was applied based on the above described SNN graph to visualize the single cell transcriptional profile in 2D space using the same number of most significant PCs. The removed TCR genes were added back to the count matrix before proceeding for downstream analyses. The cell surface protein data was normalized using the NormalizeData function with normalization.method=”CLR” (centered log ratio transformation). Briefly, for each cell, the hashtag counts were divided by the geometric mean of counts of all unique hashtags prior to log transformation.

#### Integration or merging of datasets

Experiments across different batches were integrated to mitigate batch effects using Seurat functions FindIntegrationAnchors and IntegrateData using 3000 most variable features selected across individually normalized datasets (*95*). Shared nearest neighbor graph was then calculated based on the top principal components on this integrated dataset followed by graph-based clustering and projecting the cells in a 2-dimensional space using UMAP was performed. For experiments performed in the same batch, normalized datasets were merged after filtering for common genes across datasets (*95*). Top 3000 most variable features across merged datasets were identified using SelectIntegrationFeatures and were used to calculate the most significant principal components followed by the steps mentioned in the integration workflow.

#### Differential gene expression analysis

Markers of individual clusters (or groups) were defined as the differentially expressed genes between cells belonging to each group versus every other cell using the Wilcoxon rank sum test as implemented in the function FindAllMarkers. Genes with log-fold change >0.25 and a BH-adjusted *P* value <0.05 were considered to be differentially expressed. Pairwise comparison between two clusters of interest was performed using FindMarkers function with similar cutoffs as mentioned.

#### Gene set curation and gene set enrichment analysis

Gene set enrichment scores across individual cells in clusters were quantified using AddModuleScore function in Seurat (*95*). Cytotoxicity and T_EMRA_ enriched gene sets were curated from literature (*12*), while gene lists corresponding to pairwise comparison between T cell compartments were taken from the DESeq2 comparison of our bulk transcriptomic data (data files S1 and S2). The differential expression was ordered according to the |log2 fold change| values and top 200 genes were considered for the gene sets. For *in vitro* differentiated cells, cytotoxicity module scores were used to assign cells as low-cytotoxicity (≤0), moderate-cytotoxicity (>0 ≤0.15) and high-cytotoxicity (>0.15) (Fig. 6).

#### Pseudo-time trajectory analysis

Integrated single-cell transcriptomic data from *ex vivo* CD4^+^ T cells and *in vitro* differentiated and T_H_1 polarized naïve cells were subjected to a pseudo-time trajectory analysis using Monocle3 (*72*). Seurat-based analysed integrated dataset was converted into CellDataSet objects to use them as an input in the Monocle3 framework and the UMAP generated as part of the Seurat processing pipeline was fed into this CellDataSet object. Principal graphs were built without partitions using learn_graph function. Naïve-like cells from *ex vivo* CD4 dataset, annotated based on adt_CD45RA, adt_CCR7 and the long-term memory associated T_CM_ signature score expression pattern (data file S2), were set as the root of the trajectory. Each cell was assigned a pseudo-time score and the average score across clusters were used to order the clusters based on the pseudo-time trajectory. Further, the score was used to visualize gene expression pattern changes across the trajectory using plot_genes_in_pseudotime function from the Monocle3 pipeline (*72*).

#### Single-cell TCR (V(D)J) repertoire / clonotype analysis

scTCR sequencing data was annotated using 10X Genomics’ Ensembl GRCh38 VDJ reference provided as a part of the cellranger multi pipeline and further aggregated across datasets using cellranger vdj aggr pipeline. Clonotypes in each sample were defined using the default clonotype-calling algorithm from cellranger pipeline (data file S6). Shared clonotypes were defined as clonotypes coming from different cells (≥2) for same donor. UpSet plots were used for visualizing sharing of clonotypes across clusters or cell types (*98, 99*). Circos plots, as part of the circlize package, were used for visualization of shared clonotypes maintaining the granularity of frequency, cluster and original dataset information (*100*).

### Single-cell ATAC-Seq

Single cell open chromatin analysis was performed using 10X Genomics ATAC v1.1 kit. Briefly, *ex vivo* sorted cells or *in vitro* cultured cells were washed thoroughly using 1X PBS and 55,000 viable cells were pelleted before lysing the cell membrane to isolate nuclei using the lysis buffer (10 mM Tris-HCl pH 7.4, 10 mM NaCl, 3 mM MgCl_2_, 0.1% NP40, 0.1% Tween-20, 0.01% Digitonin, 1% BSA and 1mM DTT). Lysis reaction was carried out for exact 5 minutes before wash buffer (10 mM Tris-HCl pH 7.4, 10 mM NaCl, 3 mM MgCl_2_, 0.1% Tween-20, 1% BSA and 1mM DTT) was added and nuclei were pelleted. Bulk tagmentation reaction was performed using supplied ATAC enzyme on 25,000 intact nuclei and loaded onto each lane of Chromium NextGEM Chip H. Manufacturers’ recommendations were followed to generate libraries of captured nuclei for open chromatin. Libraries were sequenced on NovaSeq 6000 or NextSeq 2000 (Illumina) as per the kit manufacturer’s recommendation.

### Single cell ATAC-Seq analysis

Demultiplexed raw ATAC library FASTQ files from each lane 10X Genomics run were mapped to the human reference genome GRCh38 using 10X Genomics’ cellranger-atac (v2.1.0) count pipeline. Cell barcode associated peak matrix (filtered for >20 <10000 bases) and corresponding fragment files were imported from cellranger run and all downstream analyses were performed using Signac v1.13.9 (*49*) unless mentioned otherwise. Barcodes were annotated as cells if at least 500 read-pairs passed read filters defined by cellranger v2.1.0. Further quality control parameters were determined based on the distribution of peak region fragment count, nucleosomal signal and TSS enrichment for each dataset and low-quality cells were filtered out: *in vitro* differentiated cells – nCount_ATAC <1000 >30000, while for *ex vivo* T_SCM_ cells – nCount_ATAC <1000 >150000; nucleosomal signal >3 and TSS enrichment <2.5 (*101*).

#### Normalization and integration

Individual peak-barcode matrices were then binarized and normalized using the implementation of the term-frequency inverse-document-frequency (TF-IDF) transformation (*49, 102*). Subsequently, singular value decomposition was run (RunSVD) on the upper quartile of accessible peaks using the most variable features selected after filtering peaks with counts >10 across all cells (FindTopFeatures(min.cutoff = 10)). Following pre-processing of individual datasets, integration was performed after projecting them into a shared low-dimensional space using reciprocal latent semantic indexing (rLSI), followed by integrating the individual dataset’s embeddings.

#### Peak calling and cluster generation

Peaks were called on this integrated dataset using MACS2 and filtered for standard chromosomal peaks (keepStandardChromosomes(pruning.mode = “coarse”)) and not aligning to ENCODE defined blacklisted regions. Fresh normalization, as mentioned above, was performed on this integrated dataset for MACS2 called and filtered peaks. Cell clusters were identified using graph-based clustering method on the shared nearest neighbor (SNN) graph using 2^nd^ through the 30^th^ dimension and the integrated LSI reduction. UMAP-based visualization was generated using the same components with a Euclidean metric. Resolution was decided empirically, for *in vitro* differentiated cells – 0.7 and for *ex vivo* T_SCM_ cells – 0.4.

#### Gene activity scores

Gene body coordinates for human genome were imported as an annotation database from TxDb.Hsapiens.UCSC.hg38.knownGene v3.16.0 from UCSC. Gene activity was then calculated using the function from Signac (GeneActivity) with regions extended 2000 bp upstream of start site (TSS) to include promoters (*49*). These gene activity scores were then log-normalized and multiplied by the median read counts per cell (nCount_reads).

#### Prediction based on scRNA-Seq analysis

Single-cell RNA-Seq experiments on same cells were performed separately and cell type annotation (based on clusters for *ex vivo* T_SCM_, Fig. 3) were transferred onto the scATAC processed data (query), based on transfer anchors from the most variable features from the scRNA dataset (reference), using canonical correlation analysis (CCA) reduction (*102*).

#### Gene signature score calculation

Gene sets curated from literature and our bulk transcriptomic analysis (data file S2) were checked for their accessibility in scATAC datasets. Accessibility scores for each gene set (module) were calculated per cell using AddChromatinModule function in Signac based on peaks linked to the genes in the set (*49*). Briefly, prior to calculation of signature scores, individual peaks are linked to the activity score for each gene in the dataset (LinkPeaks), based on Pearson’s correlation coefficient and these links are filtered for *P* value <0.05 and z-score >0.05 compared to a set of background peaks. Then the peaks associated with the genes in the given gene set (module) are filtered (GetLinkedPeaks) (data file S2; *ex vivo* T_SCM_ data: CD4-CTL enriched (Cytotoxicity signature) – 4,007 peaks, CD4-T_EMRA_ enriched (T_EMRA_ signature) – 900 peaks, CD4-T_EMRA_ vs T_N_ (200 genes) – 1,207 peaks; *in vitro* differentiation data: CD4-CTL enriched (Cytotoxicity signature) – 3,369 peaks) and deviation for the entire module is calculated per cell using chromVAR, based on the human genome BSgenome.Hsapiens.UCSC.hg38 (*103*).

#### Identification of cis-co-accessible networks (CCANs)

scATAC dataset was leveraged to identify putative cis-regulatory interactions across the gene body including promoters and enhancers to understand the regulation of gene expression across clusters. Cicero was used to predict such interactions by examining co-accessibility of peaks and was run using default parameters (run_cicero) (*54*). Cis-co-accessibility networks were then identified on co-accessible peaks using iterative cutoffs defined by the function itself (*in vitro* differentiated cells – 0.14, *ex vivo* T_SCM_ – 0.21).

#### Transcription factor motif activity analysis

Transcription factor motif activities for each cell were computed using chromVAR, based on position weight matrices (pwm) from cisbp database (*103*). The motif enrichment analysis was performed in ArchR v1.0.2 (*104*). The clustering information and the reduced UMAP coordinates for each quality control (QC) passed cell was imported from Signac into ArchR for creating the arrow object, and same parameters were followed for processing the dataset, as mentioned above. A background peak set controlling for total accessibility and GC-content was generated (addBgdPeaks), followed by addition of chromVAR motif deviations (addDeviationsMatrix) using the cisbp motif set to calculate enrichment of chromatin accessibility at different TF motif sequences in single cells. To visualize motif deviations, scores were imputed using MAGIC (*105*).

#### ChIP-Seq data integration from publicly available datasets

ChIP-Seq data for effector TFs were sourced from literature. The ChIP-seq data for T-bet was from human naïve T_H_1 polarized CD4 cells (*56*) (GEO: GSE62482 “Replicate 2”), EOMES was from hESC differentiated to endodermal fate (*57*) (GEO: GSE26097 “BoundRegions”) and FOSL2 was from human naive CD4^+^ T cells polarized to T_H_17-fate (*55*) (GEO: GSE174810 “Replicate 1”). BED files or narrowPeak files were converted to index sorted bedGraph file (-cut) which was further converted to BigWig files (bedgraphtobigwig from UCSC). Any peaks not already mapped to hg38 were lifted over, using the UCSC LiftOver tool. Histone modifications ChIP-Seq was sourced from ENCODE (https://www.encodeproject.org/). H3K27ac data was downloaded as signal *p*-value BigWig file from human CD4^+^, alpha-beta memory (CD45RO^+^) T primary cell (ENCFF884NBE from doi:10.17989/ENCSR724GUS), while H3K4me1 data was downloaded as signal *p*-value BigWig file from human CD4^+^, alpha-beta memory (CD45RO^+^) T primary cell (ENCFF334TZP from doi:10.17989/ENCSR269SSG) (*50*). The ChIP-Seq data was visualized in CoveragePlot for the genomic loci of interest with default scale.

#### Identification of motifs across genomic loci

Position Frequency Matrix (PFM) was imported from JASPAR2022 vertebrate class database as part of cellranger-arc-GRCh38-2020-A-2.0.0 reference, for TFs of interest (*73*). A motif-peak matrix was created based on the PFM for each TF for the peaks in the dataset of *ex vivo* T_SCM_ (cluster 6 and 7) and *in vitro* differentiated cells (low-, moderate- and high-cytotoxic cells) using Signac (matchMotifs(out = “positions”, p.cutoff = 0.0005), with reference genome BSgenome.Hsapiens.UCSC.hg38 (*49*). Consensus motif-containing peaks with a score cut-off >10 were then visualized using CoveragePlot for genomic regions of interest. The filtered peaks were further annotated using ChIPseeker with organism database “TxDb.Hsapiens.UCSC.hg38.knownGene” with tssRegion as -1000 to +100. The peaks which are annotated as promoters were filtered for genes in cytotoxicity gene set (data file S2) for TF of interest. UpSetR and ComplexUpset packages were used to visualize common (≥2) cytotoxicity-associated genes with promoters containing consensus motifs for TF of interest. The same data was used to create a network in Gephi to visualize the presence of consensus motifs of TF of interest in cytotoxicity-associated genes.

### Quantification and Statistical Analysis

All statistical tests on flow cytometry data were performed on percentage populations exported from FlowJo using GraphPad Prism v10.2.2. Student’s paired t-test was used to compare between two groups originating from same donors. All statistical details and population sizes are indicated in the figure legends and method details section. Population size is described in the figure legends.

## SUPPLEMENTARY MATERIALS

### Supplementary Table

**Supplementary Table S1:** List of reagents used in study.

### Supplementary Data files S1 to S7

**Supplementary Data file S1 (Microsoft Excel Format):** List of differentially expressed genes from bulk RNA-Seq analysis.

**Supplementary Data file S2 (Microsoft Excel Format):** List of gene sets used for enrichment analysis.

**Supplementary Data file S3 (Microsoft Excel Format):** Bulk TCRα and TCRβ repertoire analysis of T cell memory subsets.

**Supplementary Data file S4 (Microsoft Excel Format):** List of differentially expressed genes between clusters in single-cell transcriptomic analysis of T_SCM_ subset.

**Supplementary Data file S5 (Microsoft Excel Format):** Single-cell TCR repertoire analysis of T_SCM_ and T_EMRA_ subsets.

**Supplementary Data file S6 (Microsoft Excel Format):** List of differentially expressed genes between clusters in single-cell transcriptomic analysis of *in vitro* polarized naïve T cells at Day10 and Day20.

**Source Data (Microsoft Excel Format)**

## ACKNOWLEDGEMENTS

We thank the NII flow-cytometry core facility team (Mr. Prateek, Dr. Vikas, Mr. Khaling and Ms. Neetu) for their assistance with sorting. Our thanks to Ms. Preeti from Safdarjung hospital (SJH), Ms. Sarojini and Mr. Nandlal from IGL for assistance with sample collection at the SJH blood bank. Raji cells were a gift from Dr. Anmol Chandele. We also thank Prof. Apurva Sarin and Dr. Satyajit Rath for critically reading the manuscript.

## Funding

DBT/Wellcome Trust India Alliance Fellowship grant number IA/I/18/2/504012 (VSP).

BRIC-NII institute core funding (VSP).

DBT-JRF (Department of Biotechnology-Junior Research Fellowship) for PhD study (RK).

## AUTHOR CONTRIBUTIONS

VSP conceived the study and acquired funding for research, RK and VSP designed the research, RK, SS, ZK performed and analysed the experiments under the supervision of VSP, RK analysed the genomics data under the supervision of VSP, AS recruited, documented and collected blood samples from human donors used in this study at the blood bank, RK and VSP. wrote the manuscript, all authors read and approved the final manuscript.

## COMPETING INTERESTS

The authors declare that they have no competing interests.

## DATA and MATERIAL AVAILABILITY

Sequencing data for this study is deposited onto the Gene Expression Omnibus with series accession number GSE288989. Scripts will be made available upon request on GitHub. Any additional information required to reanalyse the data reported in this paper is available from the lead contact upon request.

## ETHICS APPROVAL STATEMENT

Ethics approval and consent to participate Approval for the use of human material for research was obtained from the institutional human ethics committees from both National Institute of Immunology, New Delhi (IHEC#119/19) and Vardhman Mahavir Medical College (VMMC) and Safdarjung Hospital (SJH), New Delhi, India (IEC/VMMC/SJH/Project/2019-08/82). The donors consented for use of the plasma and buffy coat samples for research.

## Notes

https://www.ncbi.nlm.nih.gov/geo/query/acc.cgi?acc=GSE288968

### Competing Interest Statement

The authors have declared no competing interest.

