## Supplementary figures for "A distinct subset of stem-cell memory is poised for the cytotoxicity program in CD4^+^ T cells in humans"

**Supplementary Fig. S1. Transcriptomic and TCR analysis of CD4-T cell memory compartments and CD8-$T_{EMRA}$ subset.**

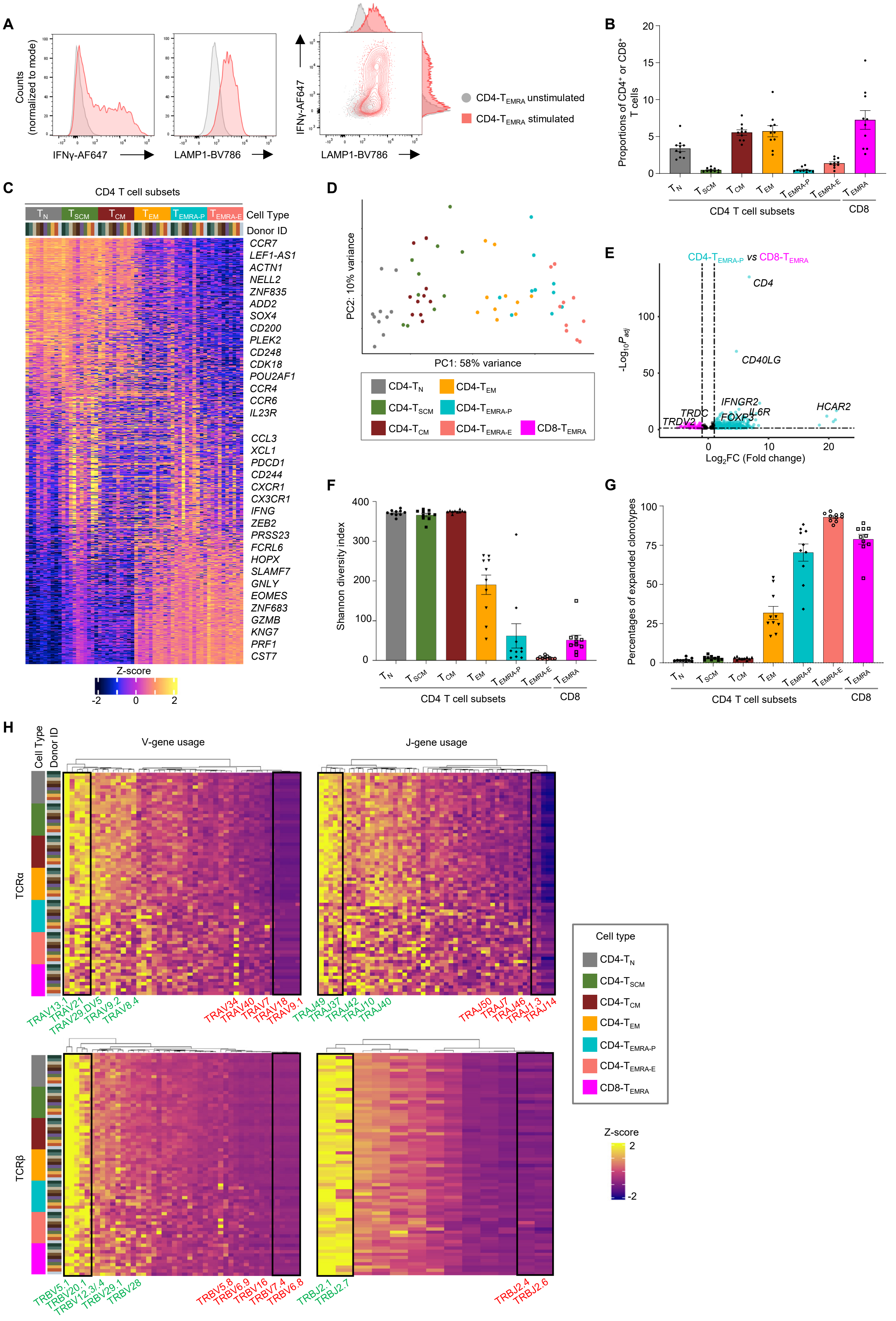

**fig. S1. Transcriptomic and TCR analysis of CD4-T cell memory compartments and CD8-$T_{EMRA}$ subset.** (A) Representative flow cytometry analysis histogram plots (left) or contour plot (right) for indicated protein expression in CD4-$T_{EMRA}$ compared between unstimulated (grey) and 6 hrs CytoStim stimulated (peach) represented using MFI for single-positives (left) or contour (right) as a function of IFNγ expression (y-axis) and LAMP1 expression (x-axis). (B) Bar graph shows the proportions of indicated memory or naïve T cell subsets within the $CD4^+$ T cell or CD8-$T_{EMRA}$ compartments in the PBMCs of 10 healthy human donors used in the RNA-Seq and TCR-seq assays. (C) Heatmap of bulk transcriptomic analysis shows the row-wise z-score of normalized counts of 1000 most variable genes across the indicated T cell memory compartments (indicated at the top) from 10 donors (each column within the cell types). (n=10). (D) Principal component analysis (PCA) plot for 1000 most variable genes across the CD4-T cell compartments. (E) Volcano plot for differentially expressed transcripts between CD4-$T_{EMRA\text{-}P}$ and CD8-$T_{EMRA}$ as a factor of -$Log_{10}$(BH-*adjusted P*) value (y-axis) and $Log_2$FC(fold change) (x-axis). Significantly differentially expressed transcripts based on DESeq2 (Benjamini-Hochberg $P_{adj}$ <0.05 and $log_2$FC >1) are colored based on cell type where they are upregulated, turquoise – CD4-$T_{EMRA\text{-}P}$ and pink – CD8-$T_{EMRA}$ (data file S1). (F-G) Bar graphs show the mean Shannon-Weiner diversity index (F) and proportions of expanded clonotypes (≥3) (G) for TCRα clonotypes in the indicated T cell compartment across 10 donors. (H) Heatmap of hierarchically clustered row-wise z-score normalized proportion of V-gene (left) and J-gene (right) usage by the recovered TCRα (top) and TCRβ (bottom). Top and bottom 5 (or 2) V- and J-genes are labelled and highlighted.

**Supplementary Fig. S2. RNA-Seq and TCR-seq analysis of CD4$^+$ T stem-cell memory ($T_{SCM}$) subset.**

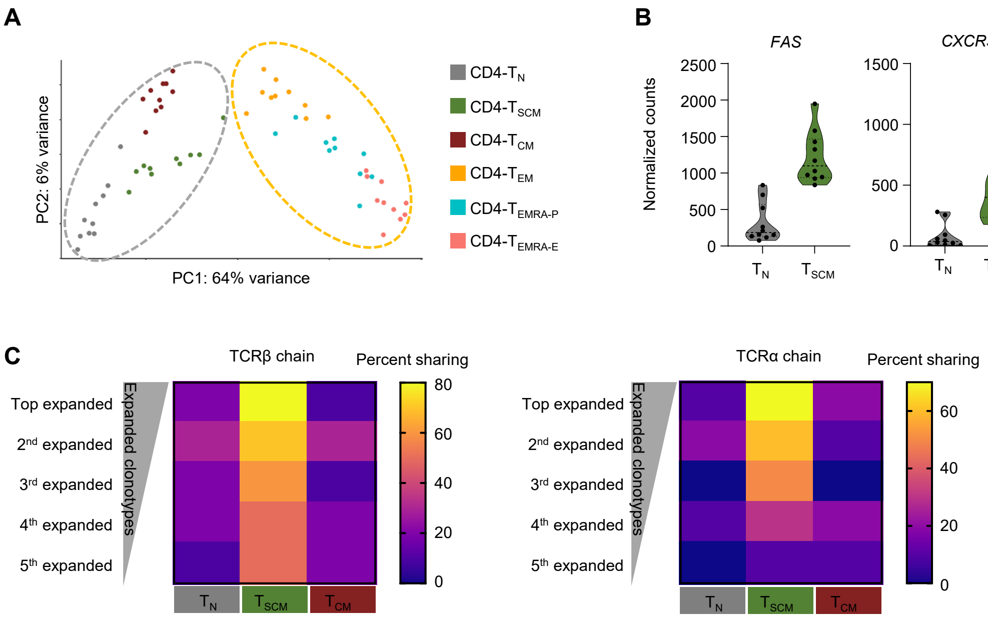

**fig. S2. RNA-Seq and TCR-seq analysis of CD4$^+$ T stem-cell memory ($T_{SCM}$) subset.** (A) Principal component analysis (PCA) plot for differentially expressed transcripts between $T_N$ and $T_{EMRA-E}$, (Benjamini-Hochberg $P_{adj}$ value <0.05 and $Log_2FC \geq 2$, 2,772 genes). (B) Violin plots show the normalized counts for indicated transcripts from RNA-Seq analysis compared between $T_{SCM}$ and $T_N$. Benjamini-Hochberg *$P_{adj}$ <0.05, **$P_{adj}$ <0.01, ***$P_{adj}$ <0.005, ****$P_{adj}$ <0.001 from DESeq2 was used for significance calculation and visualization (data file S1). (C) Heatmap of the proportion of donors (out of 10 donors) sharing the top 5 expanded TCRβ (left) and TCRα (right) clonotypes of $T_{EMRA-E}$ with $T_N$, $T_{SCM}$, or $T_{CM}$ subsets.

Supplementary Fig. S3. Identification of $T_{SCM-CTLs}$.

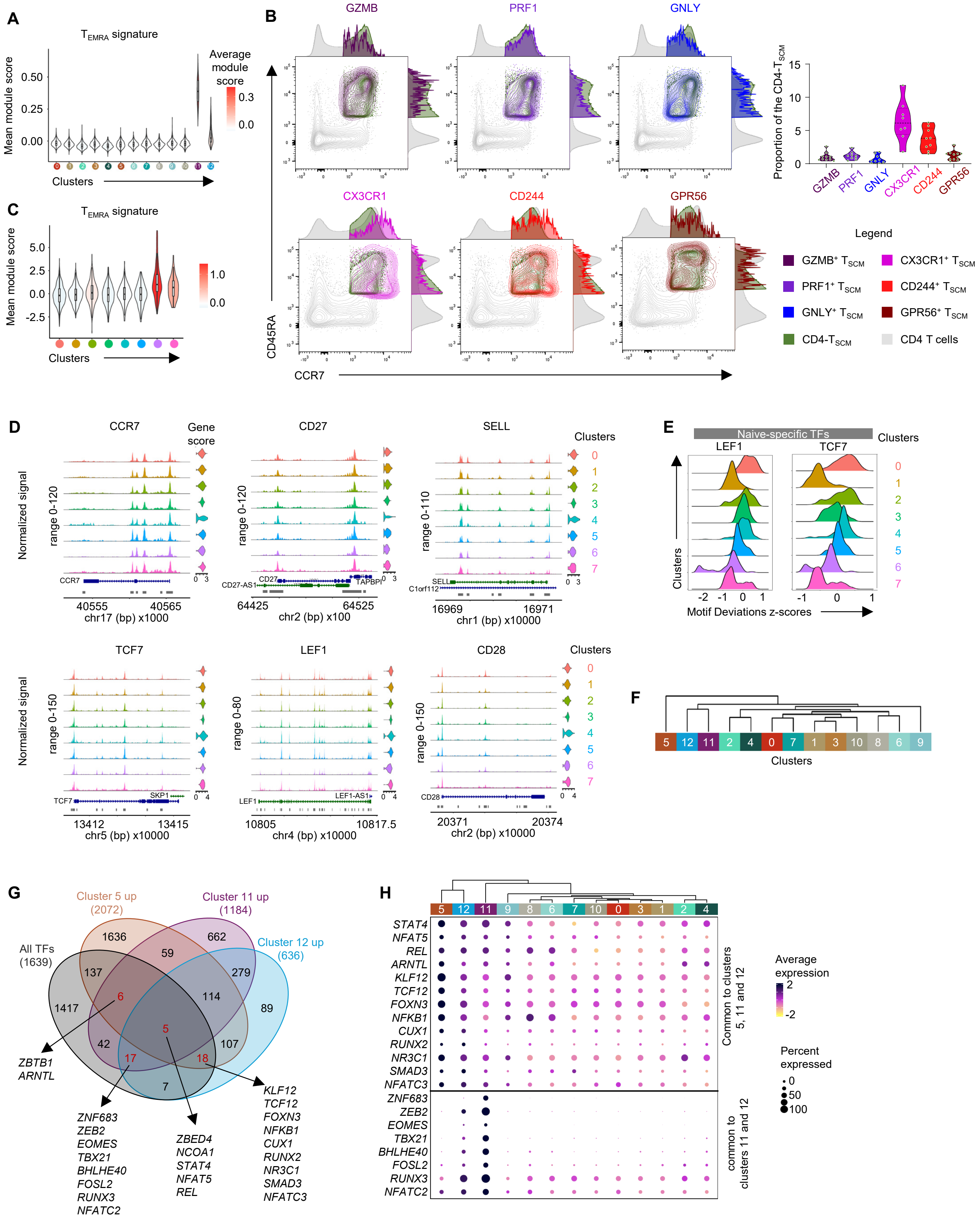

**fig. S3. Identification of $T_{SCM-CTLs}$.** (A) Violin plot for the mean signature scores for indicated gene module per cluster (x-axis) of scRNA data painted by the average score (data file S2). (B) Flow cytometry contour plots for concatenated PBMC data for 10 to 11 donors showing CCR7 (x-axis) vs CD45RA (y-axis) expression for whole CD4 T cells (light grey), overlaid by entire $T_{SCM}$ compartment (green) and individually $T_{SCM-CTLs}$ based on GZMB, PRF1, GNLY, CX3CR1, CD244 and GPR56 (color mentioned in key) expressions. Bar graph (right) shows the percentage of $T_{SCM}$ cells expressing indicated cytolytic marker protein in 10-11 donors. Each dot represents a donor. (C) Violin plot for the mean accessibility scores (y-axis) for indicated gene modules per cluster (x-axis) of scATAC data painted by the average score; CD4-$T_{EMRA}$ enriched ($T_{EMRA}$ signature) – 900 peaks (data file S2). (D) Chromatin accessibility of the indicated gene-containing genomic loci across clusters, visualized using CoveragePlot with group gene score as violin plots (right). (E) Ridge plot of chromVAR deviation (z-) scores for indicated TFs across clusters. (F) Hierarchical clustering based on the expression of 1,163 TFs from the human TF database across clusters in the scRNA-seq $T_{SCM}$ data. (G) Venn diagram shows the sharing of DE genes of clusters 5, 11 and 12 (vs rest of the clusters) with known human TFs. The DE gene list obtained from differential expression analysis, based Wilcoxon Rank Sum test with Benjamini-Hochberg $P_{adj}$ value <0.05 and $Log_2FC$ ≥0.25. Examples of overlapping genes are listed. (H) Dot plot of selected differentially expressed TFs across clusters (based on Wilcoxon Rank Sum test with Benjamini-Hochberg $P_{adj}$ value <0.05 and $Log_2FC$ ≥0.25) (data file S4). The clusters are hierarchically clustered based on the expression of shown transcripts. Color represents mean normalized and scaled expression while size represents the percentage of given transcript-expressing cells per cluster. For (A, F to H), clusters with <1% of the total cells are not shown (clusters 13 and 14 = 167 and 86 cells respectively).

**Supplementary Fig. S4. scRNA-Seq and scTCR-Seq analysis of $T_{SCM}$ and $T_{EMRA}$.**

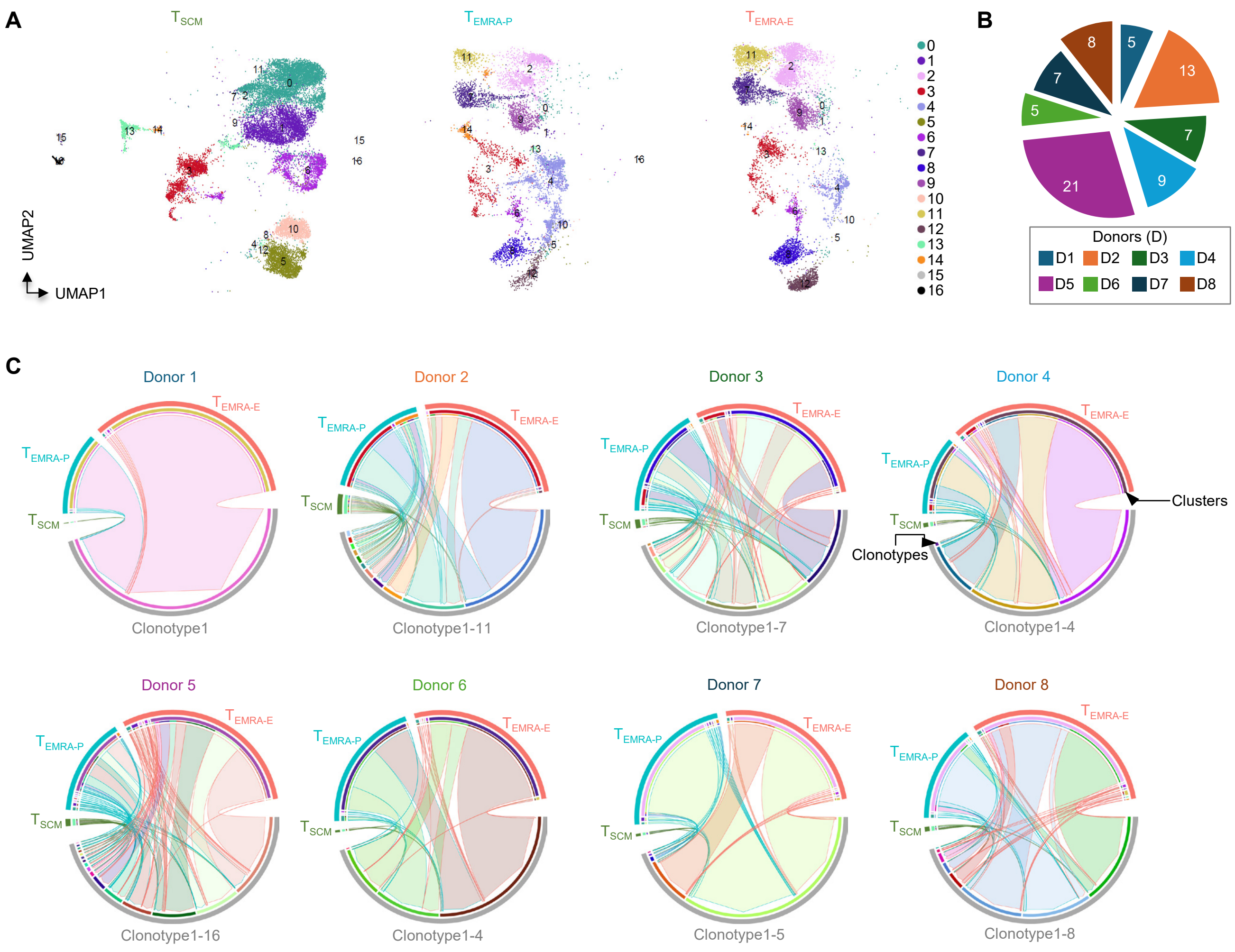

**fig. S4. scRNA-Seq and scTCR-Seq analysis of $T_{SCM}$ and $T_{EMRA}$.** (A) 2-D UMAP embedding of 33,041 cells split across $T_{SCM}$, $T_{EMRA-P}$ and $T_{EMRA-E}$. (B) Pie chart shows distribution of 75 shared clonotypes between $T_{SCM}$ and $T_{EMRA}$ (from Fig. 4E) across eight donors (D1-D8). (C) Circos plots show distribution across clusters and categories ($T_{SCM}$, $T_{EMRA-P}$ and $T_{EMRA-E}$) of 56 shared expanded clonotypes between $T_{SCM}$ and $T_{EMRA}$ cells split for each donor (D1-D8). The top half of the circos plot corresponds to the origin of the cells, with the outermost circle representing dataset origin while the circle beneath represents the originating cluster. The bottom half of the circos plot represents the clonotypes being shared with each color representing a unique clonotype. The links between the corresponding cells with their clonotypes are colored based on the clonotype with the arrowhead pointing towards the clonotype and the border of the link colored based on the origin of the cell.

**Supplementary Fig. S5. *in vitro* differentiation and polarization of naïve CD4-T ($T_N$) cells.**

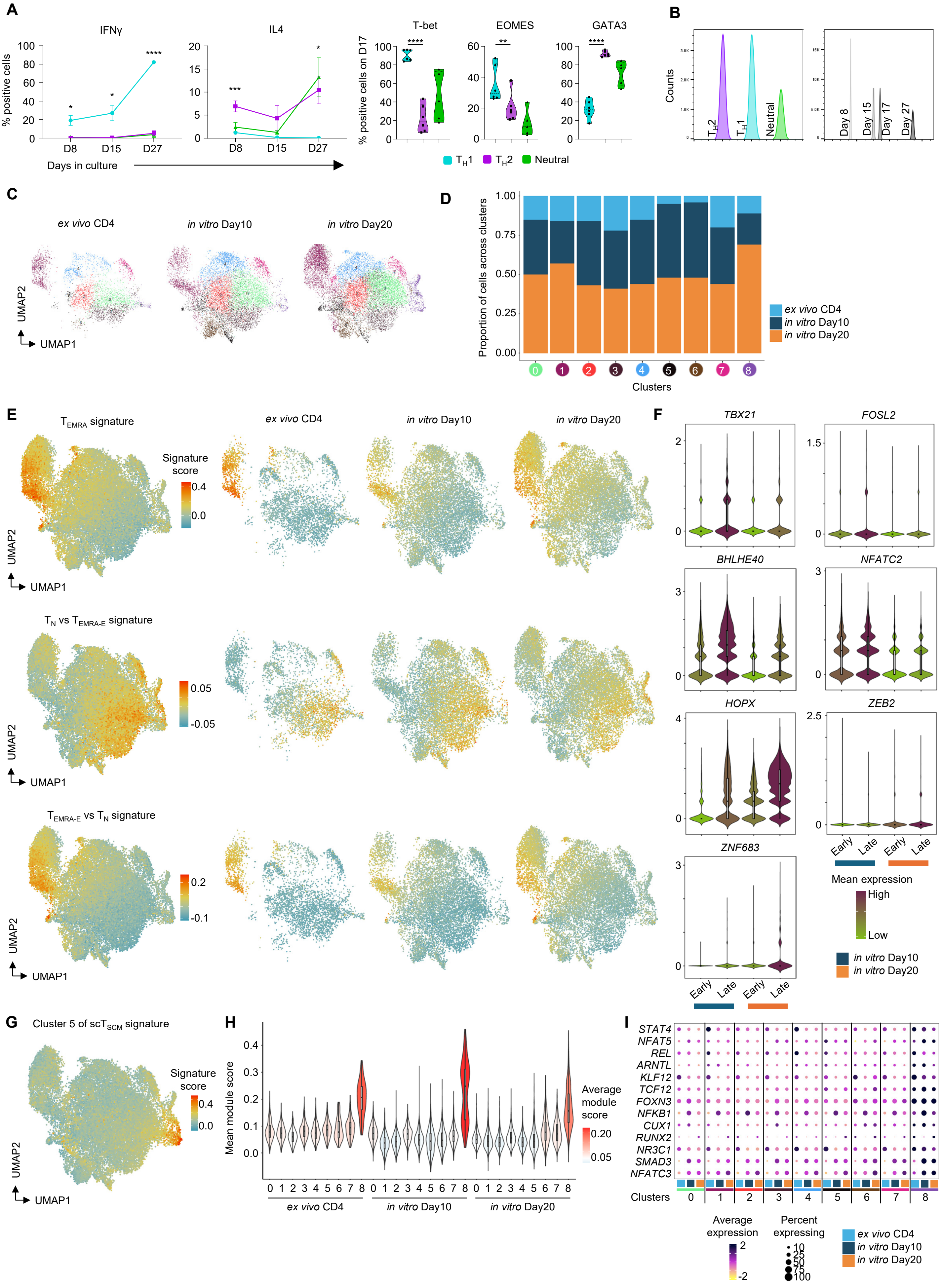

**fig. S5. *in vitro* differentiation and polarization of naïve CD4-T ($T_N$) cells.** (A) Summary flow cytometry data of indicated cytokines (IFNγ and IL4) across days comparing $T_H$1-(n=6), $T_H$2-(n=6) or neutral-(n=3) polarized cells. Error bars for each point represent mean ± SEM. Y-axis represents percentage of live-singlet gated cells expressing the mentioned protein while x-axis represents days in culture (left). Summary flow cytometry data of indicated TFs (T-bet, EOMES and GATA3) on day 17 (D17) for $T_H$1-(n=6), $T_H$2-(n=6) or neutral-(n=3) polarized cells. **P* <0.05, ***P* <0.01, ****P* <0.005, *****P* <0.001 from Student's paired two-tailed T test comparing $T_H$1 vs $T_H$2 polarizing conditions. (B) Concatenated flow cytometry data for 132,582 live-singlet gated *in vitro* differentiated and $T_H$1-, $T_H$2- and neutral- polarized cells (n=6, 6 and 3 donors respectively) from count normalized samples across donors from days 8, 15, 17 and 27 of culture, split based on polarization conditions and days, represented by count histogram. (C) Integrated 2-D UMAP embedding of 26,815 QCed-cells from *ex vivo* CD4 and *in vitro* differentiated and $T_H$1 polarized T cells, colored by clusters split by origin. (D) Stacked bar graph of proportion of cells (y-axis) from each dataset across different clusters (x-axis). (E) 2-D UMAP projection painted by indicated gene signatures either together (left) or split by original dataset (right) (data file S2). (F) Violin plots show the mean expression across groups of early (clusters 3, 0, 7 and 8) and late (clusters 2, 6, 4, 5 and 1) clusters split between D10 and D20 for indicated transcripts. (G-H) Enrichment analysis of cluster 5 (of scRNA $T_{SCM}$)-specific gene sets visualized either using 2-D UMAP projection (G) or violin plots showing module score split based on origin and clusters (H) (data file S4). (I) Dot plot of selected TFs across clusters split by origin. Color represents mean normalized and scaled expression while size represents the percentage of given transcript-expressing cells per cluster.

**Supplementary Fig. S6. $T_H1$ polarized cells undergo gradual change to acquire cytotoxicity program.**

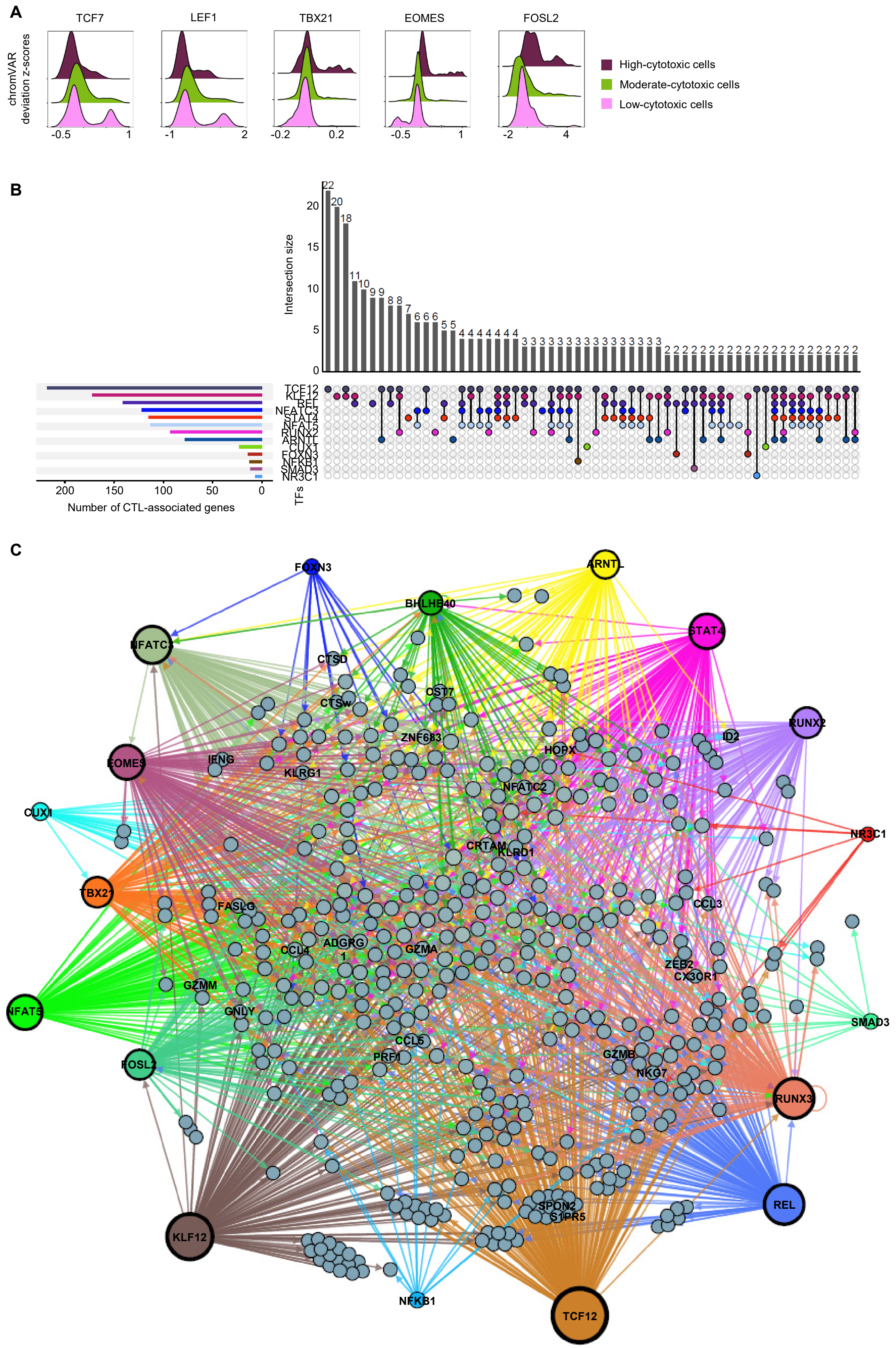

**fig. S6. $T_H1$ polarized cells undergo gradual change to acquire cytotoxicity program.** (A) Ridge plot of chromVAR deviation (z-) scores for indicated TFs in *in vitro* differentiated cells across categories (low-, moderate-, high-). (B) UpSet plot shows the frequency (y-axis) of cytotoxicity-related genes (CD4-CTL enriched (data file S2)) whose promoter has a consensus motif for the mentioned TF (x-axis) and distribution of unique or common genes (connected lines) having the consensus motif for the various TF of interest. Bar graph (left) shows the number of cytotoxicity-related genes having the consensus motif for each TF of interest at their promoter region. (C) Network plot generated using Gephi where TFs of interest have consensus motifs in the promoter of cytotoxicity-related genes (CD4-CTL enriched). Arrows indicate directionality of TF having consensus motif on target genes and colored based on TF.
